# YAP and β-catenin co-operate to drive oncogenesis in basal breast cancer

**DOI:** 10.1101/2020.06.05.115881

**Authors:** Hazel Quinn, Elle Koren, Regina Vogel, Oliver Popp, Philipp Mertins, Clemens Messerschmidt, Elisabetta Marangoni, Yaron Fuchs, Walter Birchmeier

## Abstract

Targeting cancer stem cells (CSCs) can serve as an effective approach toward limiting resistance to therapies and the development of metastases in many forms of cancer. While basal breast cancers encompass cells with CSC features, rational therapies remain poorly established. Here, we show that receptor tyrosine kinase Met signalling promotes the activity of the Hippo component YAP in basal breast cancer. Further analysis revealed enhanced YAP activity within the CSC population. Using both genetic and pharmaceutical approaches, we show that interfering with YAP activity delays basal cancer formation, prevents luminal to basal trans-differentiation and reduces CSC survival. Gene expression analysis of YAP knock-out mammary glands revealed a strong decrease in β-catenin target genes in basal breast cancer, suggesting that YAP is required for nuclear β-catenin activity. Mechanistically, we find that nuclear YAP interacts and overlaps with β-catenin and TEAD4 at common gene regulatory elements. Analysis of proteomic data from primary breast cancer patients identified a significant upregulation of the YAP activity signature in basal compared to other breast cancers, suggesting that YAP activity is limited to basal types. Our findings demonstrate that in basal breast cancers, β-catenin activity is dependent on YAP signalling and controls the CSC program. These findings suggest that targeting the YAP/TEAD4/β-catenin complex offers a potential therapeutic strategy for eradicating CSCs in basal (triple-negative) breast cancers.

## Introduction

Breast cancer is a heterogeneous human disease that has been classified into various subgroups commonly used to plan therapies^1–3^. The majority of basal breast cancers (~70%) are triple-negative for the estrogen receptor (ER), the progesterone receptor (PR) and the epidermal growth factor receptor 2 (HER2)^2,4^. Basal breast cancers are known to display high frequencies of gene alterations such as *TP53, RB* and *PTEN*, overexpression of *WNT* components, and activating mutations of receptor tyrosine kinases such as the *EGFR, MET* and others^5–10^.

Previously we generated a compound mutant mouse model that mimics the key features of human basal breast cancers^11^. It combines the activation of β-catenin, a principal downstream component of Wnt signalling^12,13^, with the expression of HGF, which activates the receptor tyrosine kinase Met^7,14^ under control of the whey acidic protein (WAP) promoter (Wnt-Met mice). *WAP* is naturally expressed in late pregnancy, thus pregnancy stimulates the rapid growth of aggressive basal mammary gland tumours in as little as two weeks postpartum^11^. Single mutants also develop tumours, but usually over a period of months. Wnt-Met tumours exhibit basal characteristics, i.e. high levels of basal markers K5, K14 and smooth muscle actin, whereas luminal cell markers K8 and K18 were low^11^. Gene expression analysis showed Wnt-Met mutant tumours grouped closely with BRCA1+/-; p53+/- basal (triple-negative) but not with luminal breast cancers^11,15^. Moreover, the expression of Wnt target genes *Lrp6, Lrp5* and *Axin* 2 were increased and several metastasis-associated genes such as *Twist1, CxCr4* and *Postn* were upregulated. Gene and protein expression studies have shown that Met is an essential protein in basal breast cancer progression and metastasis^10^. In addition, Met controls the differentiation state of Wnt-Met tumour cells, while Wnt/β-catenin controls the stem cell property of self-renewal^11^.

Basal breast cancers exhibit a heterogeneous cellular composition that includes tumour cells with stem cell-like properties^6,11,16–18^. Cancer stem cells (CSCs) have the unique capacity to initiate, maintain and replenish tumours, in contrast to other tumour epithelial cells (TECs), which makes them promising targets for cancer therapy^19,20^. However, a lack of targeted treatments generally leads to a poor five-year survival rate for patients presenting to the clinic with basal breast cancer^21^. Current therapeutic options are primarily limited to chemotherapeutic agents such as doxorubicin, which are non-specific, toxic and can reduce a patient’s overall quality of life. Although these drugs are highly effective against proliferating cells, they have little effect on CSCs due to their intrinsically low proliferation rate^2^. Increasing the survival rate for basal breast cancer patients will likely require rational therapies that target CSCs.

Here we use Wnt-Met mice to dissect the biochemical pathways of the tumours in search of components that could be targeted in therapies directed at CSCs. Our analysis of proteomic data available from Mertins et al. of primary tissue from patients revealed a high expression of the YAP signature (Yes-associated protein) in basal breast cancer, compared to HER2+, Luminal A and Luminal B tumours^22^. YAP is a critical component of the HIPPO pathway that has previously been implicated in the maintenance, survival and expansion of stem cells^23–25^. In this context, its activation is dependent on the upstream kinases LATS1/2 and MST1/2. Without these signals, the HIPPO pathway is inactive; this leads to YAP’s translocation into the nucleus, where it binds the transcriptional activator TEAD 1-4 and triggers gene transcription^26–28^. Recently a number of alternative mechanisms leading to the activation of YAP have been described, including the activation of glucocorticoid, the EGF receptor, Wnt signalling and intrinsic apoptotic components^29–32^. The aberrant activation of YAP has been shown to have a potent effect on oncogene signalling in breast cancer and many other tumours^33–36^. But this is contradicted by recent studies suggesting that YAP may also function as a tumour suppressor, or be irrelevant in breast cancer^37,38^. We speculated that these conflicting data might be due to the context-dependent role of YAP depending on tumour heterogeneity or tumour subtype.

In this study we set out to elucidate the role of YAP and its regulation in the CSCs underlying basal (triple-negative) breast cancers. Proteomic analysis identified that YAP is strongly activated by the receptor tyrosine kinase Met, contradicting a recent report suggesting that YAP is negatively regulated by RTK signalling^39^. Our genetic analysis in mice revealed that the ablation of YAP delayed tumour formation; cells with ablated YAP were located in healthy, single-layered acini with reduced proliferation. Further analyses of Wnt-Met tumours found high YAP activity in CD24^hi^CD49f^hi^ tumour-propagating cells, the generation of which was diminished upon YAP ablation. Of clinical relevance, our analysis of publicly available data confirmed a higher expression of *YAP* in the tumours of human patients with basal breast cancer than those with luminal breast cancer. In triple-negative human breast cancers, high *YAP* expression strongly correlated with reduced patient survival. Together, our findings show that YAP plays a crucial role in the initiation and maintenance of basal breast cancer and suggest that targeting YAP may represent a potential strategy for treating basal breast cancer.

## Results

### The receptor tyrosine kinase Met regulates YAP and β-catenin activity at early stages of tumorigenesis

Mice with single Wnt or Met mutations developed discernible mammary gland tumours in 30 - 40 weeks post-partum (PP), while double mutants exhibited tumours as early as two weeks PP (called Wnt-Met tumours)^11^. This led us to investigate how Wnt and Met co-operate in mammary gland tumour formation, and whether their activity is linked to that of YAP^22^. A proteomic analysis of Wnt and Wnt-Met mammary glands at early stages of tumorigenesis revealed a strong upregulation of the YAP signature^40^, indicating that YAP is activated in double but not in single mutants (Fig. 1a). We found that at early stages, Wnt-Met double mutant mammary glands contained 6-fold more YAP-positive nuclei than mice with only the Wnt mutation (Fig. 1b, c). This result was confirmed by Western blotting for active YAP (Fig. 1d, Supplementary Fig. 1a). Analysis of the phospho-proteome revealed higher levels of YAP phosphorylation at S46 and T48, suggesting that these sites play a crucial role in the phosphorylation and nuclear activation of YAP (Supplementary Fig. 1b).

**Figure 1.**
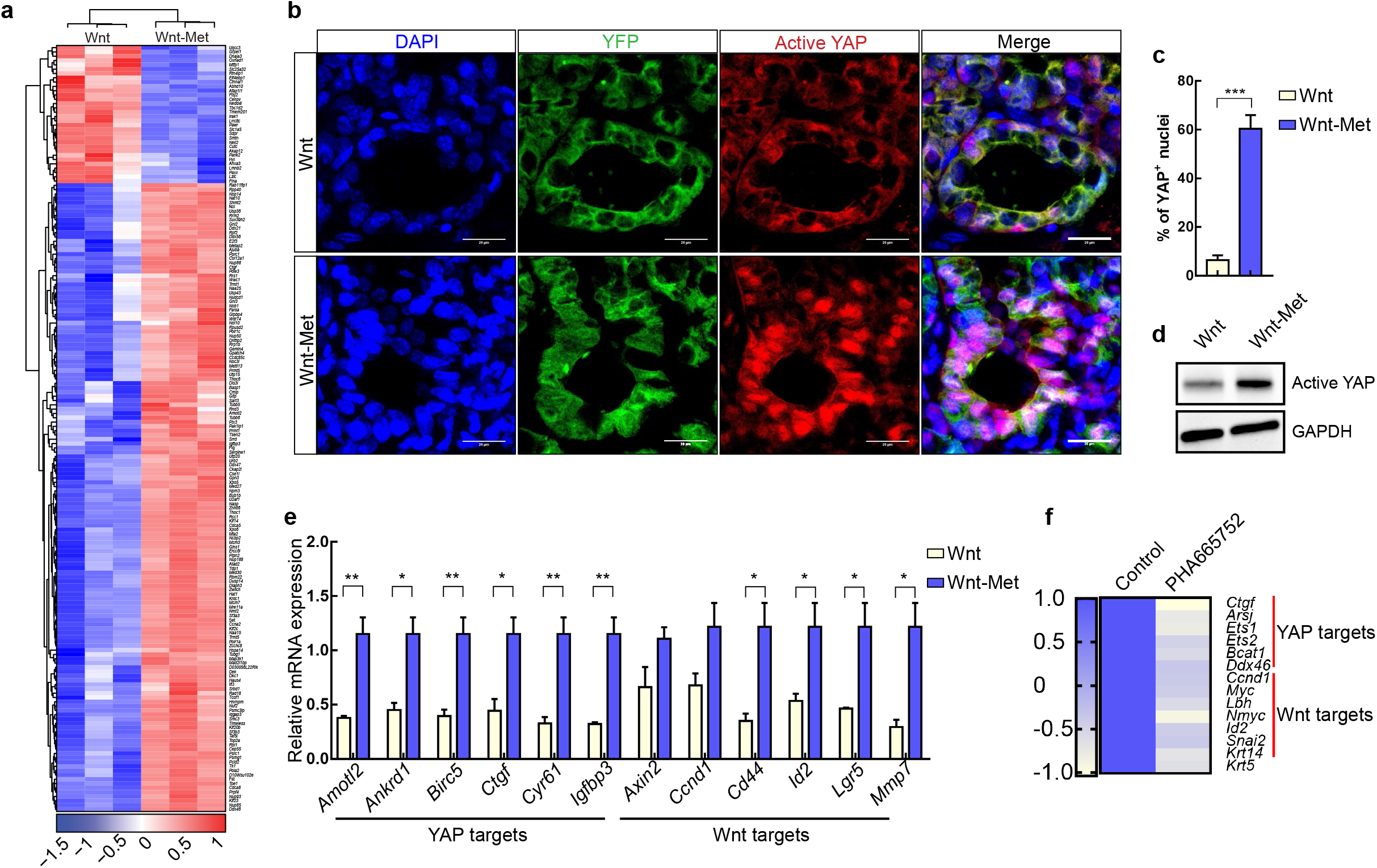
HGF-Met signalling regulates YAP activity. **a.** Heatmap of proteomic analysis showing the YAP signature (based on Zanconato et al., 2016) in WAPicre; β-cat^GOF^; ROSA26^EYFP^ (Wnt) in comparison to WAPicre; Wap-HGF; β-cat^GOF^; ROSA26^EYFP^ (Wnt-Met) at 1-week PP. The YAP signature proteins shown were significant in a two-sample moderated t-test (adj. p-value ≤ 0.05; row-scaling was applied). Values are median-MAD-normalised across all proteins and row-scaled across all samples. **b.** Immunofluorescence of YFP (green) and active YAP (red) in WAPicre; β-cat^GOF^; ROSA26^EYFP^ (Wnt) and WAPicre; Wap-HGF; β-cat^GOF^; ROSA26^EYFP^ (Wnt-Met) at 1-week PP, scale bar, 20μm. **c.** Quantification of YAP-positive nuclei (YAP^+^ nuclei/number of nuclei per field x100) in Wnt vs. Wnt-Met tissues. Data shown are mean ±SEM, n=3 biological replicates, ***p<0.001, by Student’s t test. **d.** Western blot of active YAP in Wnt and Wnt-Met tissues at 1 week PP. **e.** RT-qPCR of YAP and Wnt target genes in Wnt and Wnt-Met tissues at 1 week PP. Data shown are mean ±SEM, n=3 biological replicates, *p<0.05, **p<0.01, by Student’s t-test. **f.** Heatmap of YAP and Wnt target genes of PHA665752-treated mammospheres^11^.

We also examined whether Met signalling is responsible for the nuclear translocation of β-catenin. Double mutants exhibited a 10-fold increase of nuclear β-catenin compared to the mammary glands of mice with only the Wnt mutation (Supplementary Fig. 1c and d). The expression of YAP and Wnt target genes including *Birc5, Ctgf, Cd44* and *Lgr5* was significantly higher in the mammary glands of Wnt-Met mice (Fig. 1e)^41,42^. We also examined the expression of YAP and Wnt target genes in Wnt-Met mammospheres^43,44^ treated with the Met inhibitor PHA665752^11,40,45^. Remarkably, this revealed significantly lower levels in the expression of *Ctgf, Ccnd1 (CyclinD1)* and other genes, confirmed by qPCR (Fig. 1f, Supplementary Fig. 1e)^40,45^. These data show that the RTK Met is required for YAP and β-catenin nuclear translocation in the mammary glands of Wnt-Met mice.

### Genetic and pharmacological evidence that basal mammary gland tumours are dependent on YAP activity

Our finding that YAP is highly activated in basal breast cancer led us to examine its functional role through genetic and pharmacological interference. We crossed floxed YAP alleles^46^ into Wnt-Met mice. Upon stimulation via pregnancy, this led to homozygous YAP ablation in Wnt-Met mice (denoted Wnt-Met-YAP^KO^) and produced an increase in the number of tumour-free mice from 11 days in controls (Wnt-Met and heterozygous YAP ablation) to 17days (Fig. 2a). The average weight of Wnt-Met-YAP^KO^ mammary gland tumours was also decreased (Fig. 2b). We confirmed *YAP* ablation by qPCR (Fig. 2c). Remarkably, H&E staining and immunohistochemistry of Wnt-Met-YAP^KO^ mammary glands revealed large alveoli which were mostly YAP-free and exhibited healthy one-layered acini. Their proliferation was lower, as determined by Ki-67 staining (Fig. 2d, lower panel, quantification in Fig. 2e). In contrast, Wnt-Met-YAP^Ctrl^ mammary glands showed large tumorous areas which were strongly YAP-stained, without large, empty alveoli (Fig. 2d, upper panel). Fluorescent images of YFP confirmed the successful Cre-recombination of the mammary epithelial cells; YFP-positive cells were found in single layered acini in Wnt-Met-YAP^KO^ glands, in contrast to filled tumours in the controls (Fig. 2f).

**Figure 2.**
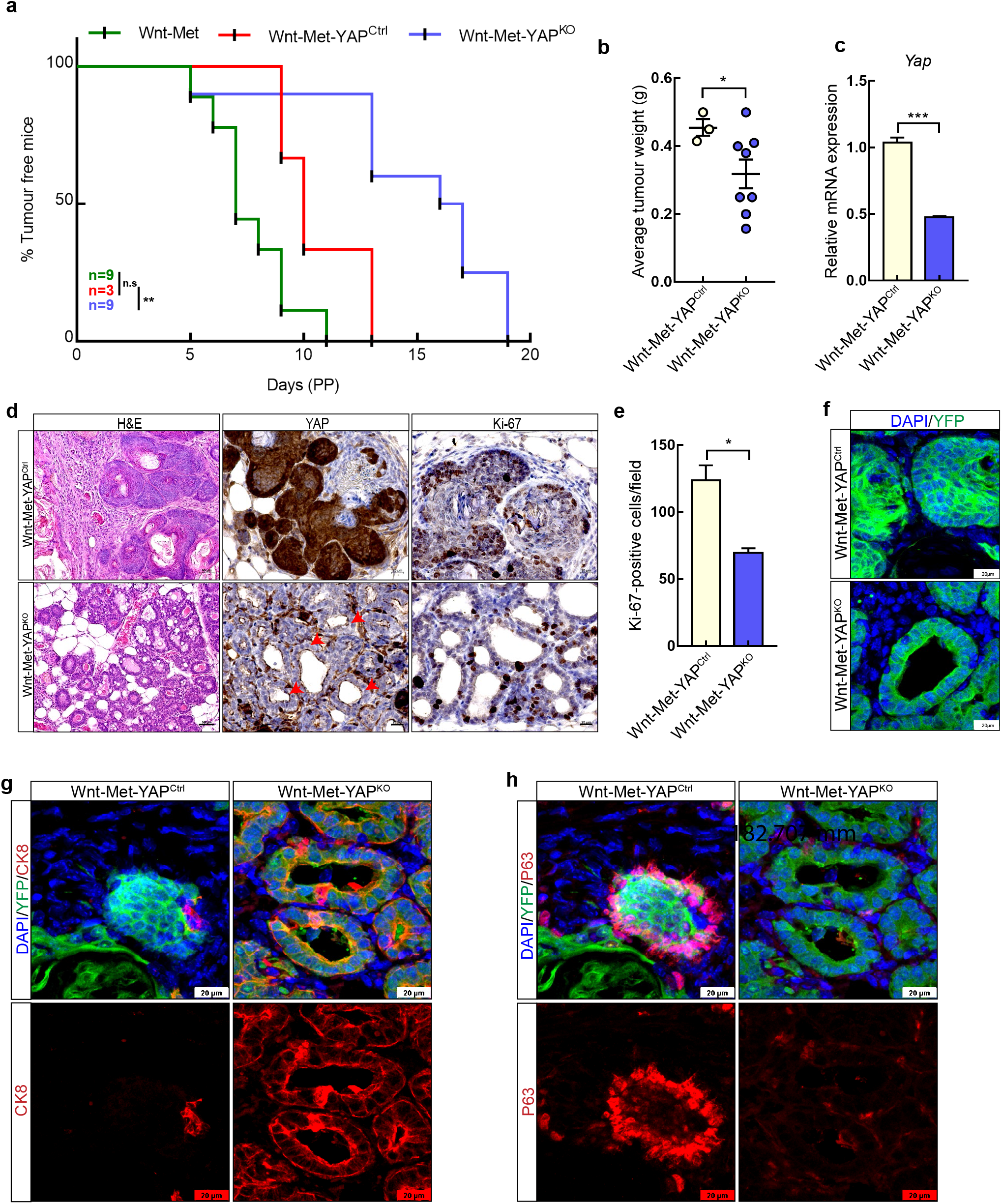
Genetic ablation of YAP delays Wnt-Met tumour formation. **a.** Graph showing tumour-free mice in Wnt-Met, Wnt-Met-YAP^Ctrl^ and Wnt-Met-YAP^KO^, **p<0.001, by Gehan-Breslow-Wilcoxon test. **b.** Average tumour weight of Wnt-Met-YAP^Ctrl^ and Wnt-Met-YAP^KO^ mice. Data shown are mean ±SEM, *p<0.05, by Student’s t-test. **c.** RT-qPCR of *Yap* mRNA expression in Wnt-Met-YAP^Ctrl^ and Wnt-Met-YAP^KO^ tumours. Data shown are mean ±SEM, n=3 biological replicates, ***p<0.001, by Student’s t test. **d.** Comparison of Wnt-Met-YAP^Ctrl^ (upper pictures) and Wnt-Met-YAP^KO^ (lower pictures) mammary glands at 2.4 weeks PP. H&E staining (left panel), immunohistochemistry of YAP (middle panel) and Ki-67 (right panel), red arrowheads mark single cell-layered healthy YAP-free epithelia, scale bar, 50μm and 20μm. **e.** Quantification of Ki-67-positive cells. Data shown are mean ±SEM, n=3 biological replicates, *p<0.05, by Student’s t test. **f.** Confocal images of YFP (green) in Wnt-Met-YAP^Ctrl^ and Wnt-Met-YAP^KO^ mammary glands at 2.4weeks PP, scale bar, 20μm. **g.** Confocal images of YFP (green) and CK8 (red) in Wnt-Met-YAP^Ctrl^ and Wnt-Met-YAP^KO^ mammary glands at 2.4 weeks PP, scale bar, 20μm. **h.** Confocal images of YFP (green) and P63 (red) in Wnt-Met-YAP^Ctrl^ and Wnt-Met-YAP^KO^ mammary glands at 2.4 weeks PP, scale bar, 20μm.

We treated Wnt-Met mice systemically with two inhibitors of YAP activity. Simvastatin (SIM) inhibits YAP via the mevalonate-Rho kinase pathway^47^, and verteporfin (VP) inhibits the interaction of YAP with its transcriptional activator TEAD^48^. Both inhibitors reduced tumour volumes and weight (Supplementary Fig. 2a-d). H&E and immunohistochemistry of SIM and VP-treated tumours also revealed small, empty alveoli and a strong reduction in Ki-67-positive cells compared to controls (Supplementary Fig. 2e-j). In combination, these data demonstrate that YAP is crucial for tumour initiation in Wnt-Met tumours.

Met signalling induces the trans-differentiation of luminal mammary gland cells into basal cells^11,49^. To examine whether Met-induced trans-differentiation also depends on YAP, we examined Wnt-Met-YAP^Ctrl^ and Wnt-Met-YAP^KO^ mammary glands for the expression of the luminal cell maker CK8 and the basal cell marker P63. We observed strong expression of P63 in control glands, which exhibited a minimal expression of CK8 (Fig. 2g, h, left panels). Strikingly in contrast, Wnt-Met-YAP^KO^ mammary glands showed negligible P63-positive cells but a high number of CK8-positive cells (Fig. 2g, h, right panel). As further confirmation, in Wnt-Met tumours we found a strong expression of CK14 and a low expression of CK8, a pattern which was reversed in SIM-treated tumours (Supplementary Fig. 2k). Overall, these data provide genetic and pharmacological evidence that YAP is required for the acquisition of basal characteristics in Wnt-Met tumours.

### YAP is active in the cancer stem cells of basal mouse mammary gland tumours

We have previously shown that Wnt-Met tumours induced by WAP-Cre harbour CD24^hi^ and CD49f^hi^/CD29^hi^ cancer stem cells that produce aggressive basal cancers when transplanted into immune-deficient mice^11^. We now examined the contributions of YAP to Wnt-Met CSCs. Using an antibody specific for the active state of YAP, we found high levels of nuclear YAP on the outer rim of Wnt-Met tumour acini, which correlated to the increase of expression of specific stem cell genes such as CD44 and Sox10 (Fig. 3a, Supplementary Fig. 3a, b)^18,16,50–52^. We confirmed high levels of YAP by Western blotting and mRNA expression (Fig. 3b, c). Moreover, the expression of target genes of Wnt/β-catenin and YAP including *Axin2*^53^, *Ccnd1*, *Ctgf*^40^ and *Igfbp3* increased (Fig. 3d, e).

**Figure 3.**
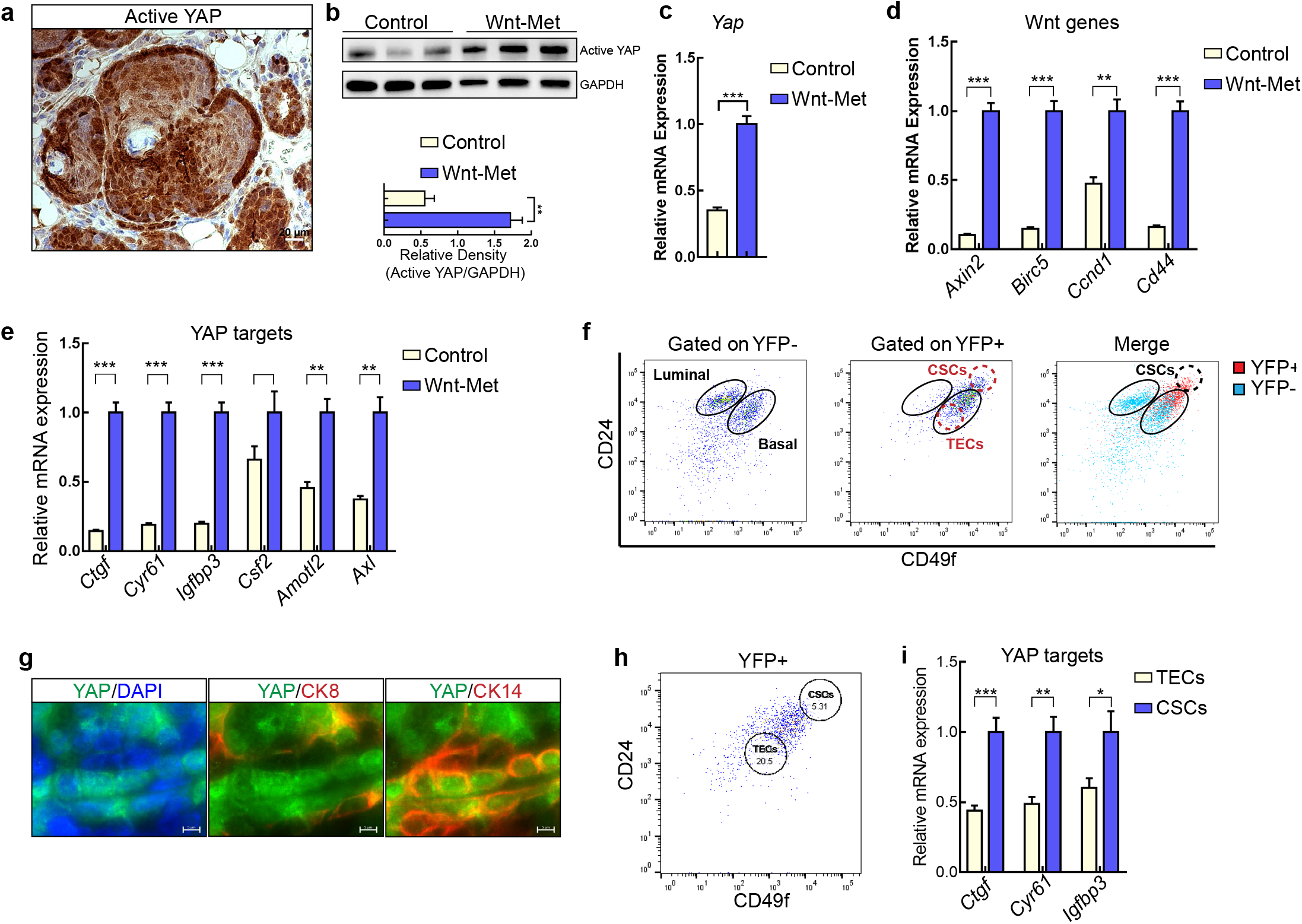
YAP is activated in the CSCs of Wnt-Met-driven mammary gland tumours. **a.** Immunohistochemistry of active YAP in Wnt-Met tumours at 2-weeks post-partum (PP), scale bar, 20μm. **b.** Western blot of control and Wnt-Met tumours at 2 weeks PP showing active YAP and GAPDH (above), and quantification of active YAP expression in control and Wnt-Met tumours 2-weeks PP (below). Data shown as mean ± SEM, n=3 biological replicates **p<0.01 by Student’s t test. **c.** RT-qPCR of *Yap* expression in control and Wnt-Met tumours 2 weeks PP. Data shown as mean ± SEM, n=3 biological replicates ***p<0.001 by Student’s t test. **d.** RT-qPCR of Wnt target genes in control and Wnt-Met tumours 2 weeks PP. Data shown as mean ± SEM, n=3 biological replicates **p<0.01, ***p<0.001 by Student’s t-test. **e.** RT-qPCR of YAP target genes in control and Wnt-Met tumours at 2 weeks PP. Data shown as mean ± SEM, n=3 biological replicates **p<0.01, ***p<0.001, by Student’s t-test. **f.** FACS for YFP, CD24 and CD49f. **g.** Immunofluorescence of YAP (green), CK8 (red) and CK14 (red), scale bar, 5μm. **h.** FACS showing isolation of cancer stem cells (CSCs) and tumour epithelial cells (TECs). **i.** RT-qPCR of YAP target genes in TECs and CSCs. Data shown are mean ±SEM, n=5 biological replicates, *p<0.05, **p<0.01, ***p<0.001, by Student’s t-test.

High levels of YAP activity have been reported in the stem cell compartments of many tissues^54,55^. CD24^hi^CD49f^hi^ cancer stem cells were found in the basal cell population of mammary gland tumours but not the luminal, as shown by FACS (Fig. 3f, Supplementary Fig. 3c). In agreement with this, CSCs have been reported among the basal cells marked by cytokeratin 14 (CK14)^11,56^. We found that YAP was expressed in the nucleus of basal, CK14-positive cells but not in those that were luminal and CK8-positive (Fig. 3g). Moreover, YAP target genes were more highly expressed in CSCs than other tumour epithelial cells (TECs) (Fig. 3h, i). These data show that Wnt-Met signalling generates basal tumours with high YAP activity in CSCs of the mammary gland.

### YAP regulates growth and initiation of Wnt-Met cancer stem cells

To study the function of YAP in CSCs, we first established stem cell-enriched control and Wnt-Met spheres in mammary gland stem cell-promoting medium^57,58^ (Supplementary Fig. 4a). Control spheres (WAP-Cre; ROSA26^EYFP^) formed structures with single-layered epithelia with empty lumens, in contrast to the large, filled structures produced by Wnt-Met spheres (Fig 4a, Supplementary Fig. 4b). FACS for YFP showed that Wnt-Met spheres contained 1.4-fold (57% to 76%) more YFP-positive cells than controls (Supplementary Fig. 4c). They also exhibited a higher expression of transgenes, as confirmed by qPCR for *Wap, Hgf* and *Lgr5* (Supplementary Fig. 4d). Immunofluorescence demonstrated a decrease in the luminal marker CK8 and an increase in the basal marker CK14 (Supplementary Fig. 4e). Moreover, a strong nuclear β-catenin signal was observed in the basal cells located at the outer rim of these spheres (Fig. 4b). We next isolated cells by FACS and seeded equal cell numbers to generate stem cell-enriched spheres. The CSC population underwent an 8-fold higher outgrowth than TECs (Supplementary Fig. 4f). These results show that sphere conditions enrich and expand CSCs in the Wnt-Met mammary gland tumours. We then identified nuclear YAP in tumour-derived spheres through immunofluorescence (Supplementary Fig. 4g) and quantified the expression of YAP target genes using qPCR. We found a 2.5-fold increase in *Cyr61*^40^ and 3.7-fold increase in *Igfbp3*^40^(Supplementary Fig. 4h).

**Figure 4.**
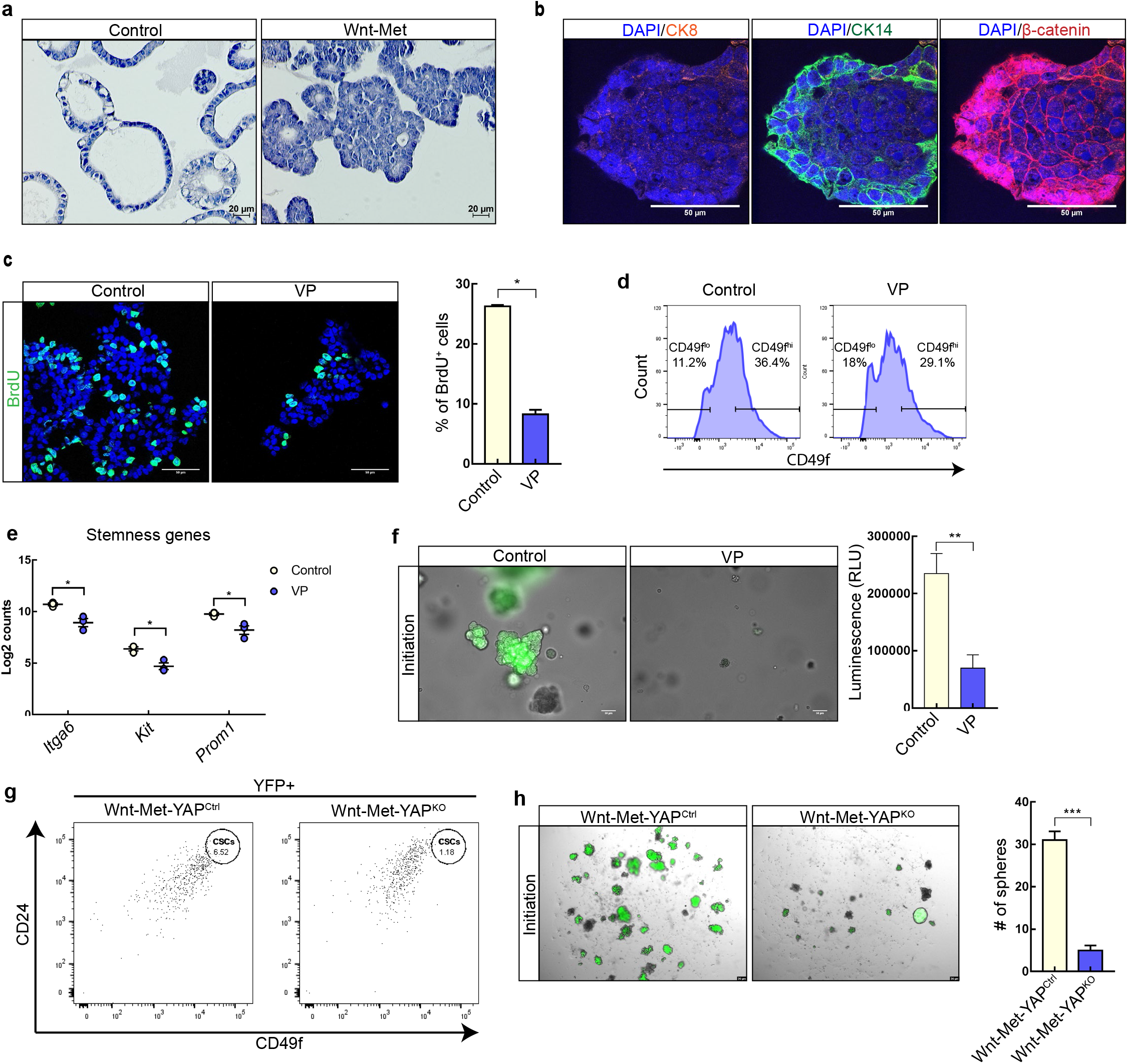
YAP regulates stem cell properties. **a.** H&E staining of sectioned stem cell-enriched spheres generated from control and Wnt-Met tumours, scale bar, 20μm. **b.** Confocal microscopy of Wnt-Met stem cell-enriched spheres for CK8 (orange), CK14 (green) and β-catenin (red), scale bar, 50μm. **c.** Confocal images of BrdU (green) of stem-enriched spheres treated with vehicle control (left) or 2μM verteporfin (right), quantification on the right. Data shown are mean ±SEM, n=3 biological replicates, *p<0.05, by Student’s t test. **d.** FACS histogram showing the number of CD49f^hi^ cells in control or VP-treated stem cell-enriched spheres, 1.25-fold difference. **e.** Log2 counts of *Itga6, Kit*, and *Prom1* mRNAs from control or VP-treated stem cell-enriched spheres. Data shown are mean ±SEM, n=3 biological replicates, *p<0.05, by Student’s t-test. **f.** Bright-field and fluorescent images of Wnt-Met stem cell-enriched spheres treated from day one with vehicle control (left) or 2μM VP (right). Quantification is on the right, scale bar, 50μm. Data shown are mean ±SEM, n=3 biological replicates, **p<0.01, by Student’s t test. **g.** FACS plot showing the percentage of YFP^+^, CD24^hi^, CD49f^hi^ cells in Wnt-Met-YAP^Ctrl^ and Wnt-Met-YAP^KO^ mammary glands. **h.** Bright-field and fluorescent images of Wnt-Met stem cell-enriched spheres generated from Wnt-Met-YAP^Ctrl^ and Wnt-Met-YAP^KO^ tissue, scale bar, 20μm. Quantification of sphere numbers is on the right. Data shown are mean ±SEM, n=3 biological replicates, ***p<0.001, by Student’s t test.

To understand the functional role of YAP, we inhibited its activity using SIM and VP. These treatments reduced proliferation of Wnt-Met spheres, as determined by BrdU and Ki-67 immunofluorescence (Fig. 4c, quantification on the right, Supplementary Fig. 4i). The inhibition of YAP in Wnt-Met spheres reduced the CSCs marked by CD49f 1.25-fold (36% to 29%) as measured by FACS (Fig. 4d). An expression analysis for the stem cell-associated genes *Itga6, Kit*^59^ and *Prom1*^60^ further supported the inhibition of CSCs (Fig. 4e). Moreover, sphere initiation identified by the number of spheres was repressed 2.8-fold by VP (Fig. 4f, quantification on the right). Since VP may have off-target affects^61^ we also treated cells with YAP-directed shRNA and siRNA which resulted in a 4-fold decrease of the size of spheres generated by Wnt-Met cells and a 2.5-fold decrease in the number of spheres by MDA-231 cells (Supplementary Fig. 4j-m). FACS analysis revealed a 5.5-fold decrease in the content of CD49f^hi^CD24^hi^ CSCs^11^ in Wnt-Met-YAP^KO^ mammary glands compared to controls (Fig. 4g). Furthermore, spheres generated from Wnt-Met-YAP^KO^ mammary glands generated 6-fold fewer spheres than controls (Fig. 4h, Supplementary Fig. 4n). This demonstrates that YAP activity is essential for the growth, initiation and self-renewal of Wnt-Met CSCs.

### YAP is required for β-catenin activity in Wnt-Met mammary gland tumours

We carried out Nanostring gene expression analysis^62^ of control and Wnt-Met-YAP^KO^ mammary glands. Wnt-Met-YAP^KO^ mammary glands exhibited a strong decrease in YAP and β-catenin activity (Fig. 5a). This tissue exhibited a downregulation of target genes such as *Ctgf* and *Igfpb3* (YAP targets) and *Axin2* and *BMP4* (β-catenin targets); the same was found for VP-treated spheres (Fig. 5b, c, Supplementary Fig. 5a). Quantification showed that β-catenin-positive nuclei were decreased 11-fold by the ablation of YAP (Fig. 5d, quantification on the right) and a strong decrease by pharmacological inhibition of YAP (Supplementary Fig. 5b).

**Figure 5.**
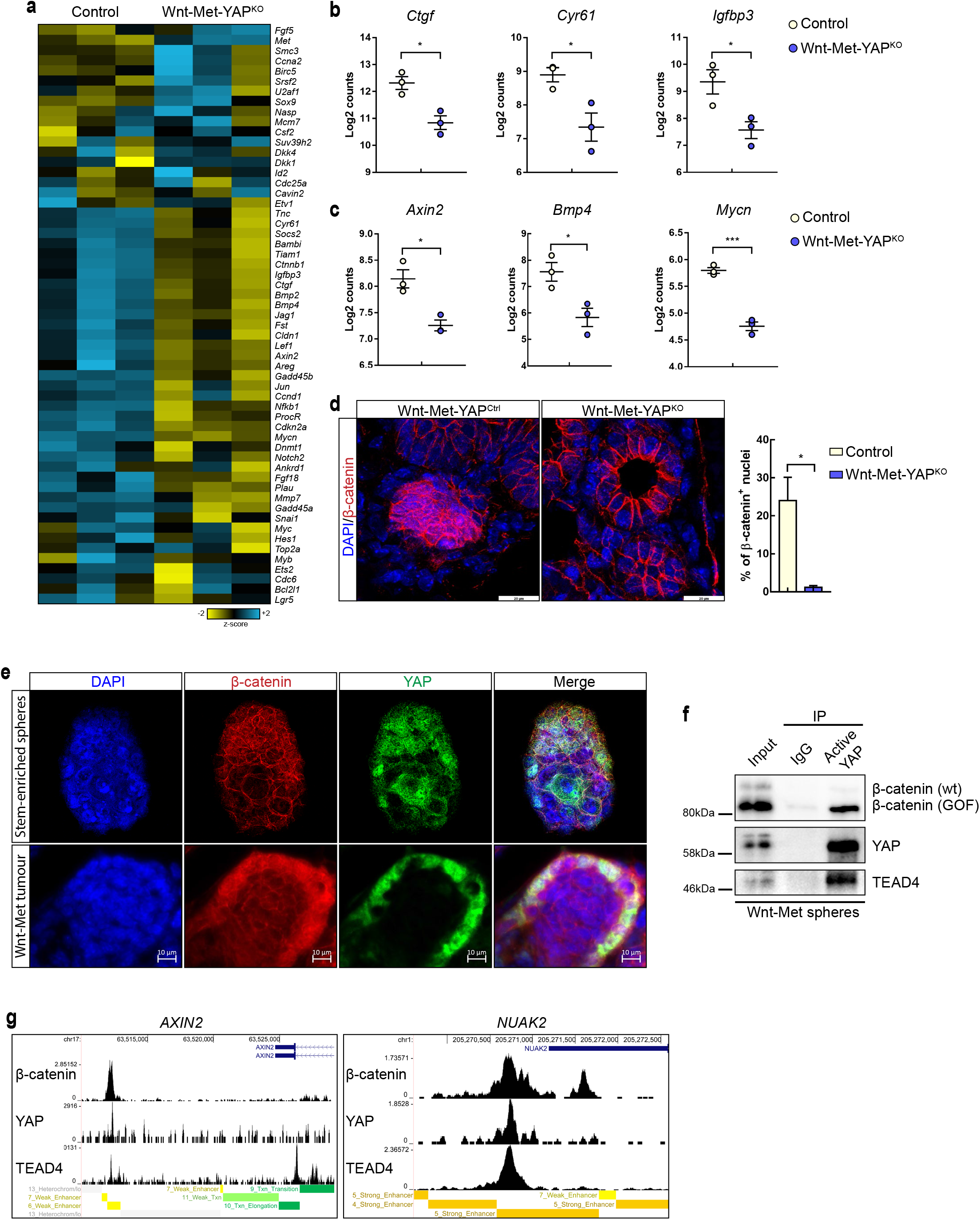
β-catenin activity is YAP dependent in mammary gland tumours. **a.** Heatmap showing differential gene expression of YAP and β-catenin target genes in Wnt-Met-YAP^Ctrl^ and Wnt-Met-YAP^KO^ mice. **b.** Log2 counts of the YAP target genes *Ctgf, Cyr61* and *Igfbp3*. **c.** Log2 counts of the β-catenin target genes *Axin2, Bmp4* and *Mycn*. Data are mean ±SEM, n=3 biological replicates, *p<0.05, ***p<0.001, by Student’s t test. **d.** Confocal images of β-catenin (red) in Wnt-Met-YAP^Ctrl^ and Wnt-Met-YAP^KO^ mice. Quantification of nuclear β-catenin on the right. Data are mean ±SEM, n=3 biological replicates, *p<0.05, by Student’s t test. **e.** Confocal images of β-catenin (red) and active YAP (green) in stem cell-enriched spheres (upper) and Wnt-Met tumours (lower), scale bar, 10μm. **f.** Western blot showing co-immunoprecipitation of active YAP with β-catenin and TEAD4 in Wnt-Met spheres. **g.** ChIP-seq tracks for YAP, β-catenin, TEAD4 showing overlapping signals at enhancer regions.

In the absence of Wnt signalling, an interaction of YAP and β-catenin has been implicated in the β-catenin destruction complex^30^. Here, immunofluorescence revealed that YAP and β-catenin are co-localized in the nucleus of Wnt-Met spheres and tumours (Fig. 5e). Active YAP co-immuno-precipitated with β-catenin^GOF^ and Tead4 in Wnt-Met spheres, but to a lesser extent with wildtype β-catenin (Fig. 5f).

YAP has been reported to be required for the development of β-catenin driven tumours^63^. Since YAP inhibition resulted in a decrease in the expression of a number of Wnt target genes, we hypothesized that YAP/TEAD might bind with β-catenin at promoter and/or enhancer regions of common target genes to regulate their expression in CSCs. To test this, we obtained YAP1, TEAD4 and CTNNB1 (β-catenin) chIP-seq data from chIP-Atlas^64^. Since YAP has been reported to preferentially bind enhancer regions rather than promoters^65^, we investigated whether YAP/TEAD4/CTNNB1 could occupy an enhancer region of the Wnt target gene *AXIN2*^53^. We identified a distal enhancer of *AXIN2* and a strong enhancer of *NUAK2* (a newly identified YAP target gene^66^) containing overlapping CTNNB1, TEAD4 and YAP1 chIP-seq peaks (Fig. 5g). Overall, these data show that TEAD4, CTNNB1 and YAP1 interact and can co-locate at gene promoters and enhancers of both Wnt and YAP target genes.

### YAP is active in human basal breast cancers and predicts patient survival

The analysis of human primary patient proteomic data available from Mertins et al.^22^ revealed high expression of the YAP signature^40^ in basal breast cancer, in contrast to other subgroups of breast cancers (Fig. 6a). qPCR of YAP target genes *CTGF, IGFBP3* and *ANKRD1*^40^ in spheres of human basal breast cancer cell lines BT549, MDA231 and SUM1315 confirmed elevated YAP activity compared to the luminal cell lines MCF7, BT474 and T47D (Fig. 6b, Supplementary Fig. 6a). Online datasets available from Gene expression-based Outcome for Breast cancer Online (GoBo)^67^ confirmed a significantly higher expression of YAP in tumours of human patients with basal breast cancer than in tumours of patients with luminal breast cancer (Supplementary Fig. 6b).

**Figure 6.**
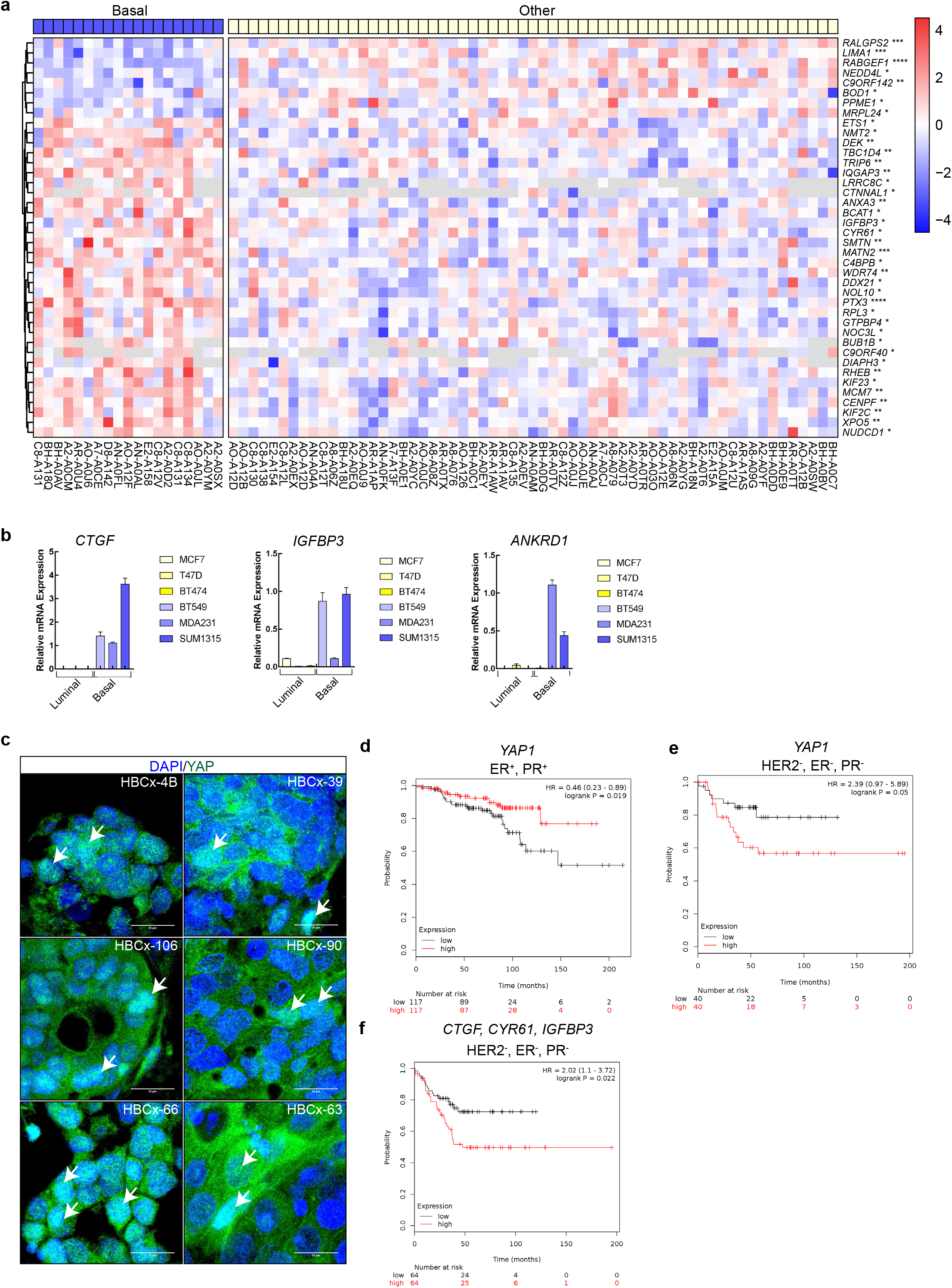
YAP is active in human basal breast tumours and predicts patient outcome in a subtype-dependent manner. **a.** Heatmap of proteomics data from Mertins et al.^40^. The provided CPTAC dataset was filtered for samples that passed the QC criteria. A two-sample moderated t-test was applied between samples that have been assigned to basal vs. all other subtypes combined. Selected significant proteins from Zanconato et al.^40^ with an adjusted p-value from Benjamini-Hochberg correction < 0.05 from the lists are displayed. The heatmap uses median-MAD-normalised input data across all proteins and row-scaling across all samples. Hierarchical clustering by Euclidian distance was applied to rows, while missing values are indicated by grey colour. Significance as indicated: *adj. p-value < 0.05, **adj. p-value < 0.01, ***adj. p-value < 0.001, ****adj. p-value < 0.0001. **b.** RT-qPCR analysis of YAP target genes in luminal (yellow) and basal (blue) cell lines. **c.** Confocal immunofluorescence of nuclear YAP in basal PDX models, scale bar, 20μm. **d.** Kaplan-Meier plot showing disease-free survival (DFS) of ER-positive and PR-positive patients with high (red) and low (black) YAP expression. **e.** Kaplan-Meier plot showing disease-free survival of triple-negative breast cancer patients with high (red) and low (black) YAP expression. **f.** Kaplan-Meier plot showing the DFS of triple-negative breast cancer patients with high (red) and low (black) expression of YAP target genes.

Immunofluorescence analysis of active YAP in six basal breast cancer PDX models^68^ exhibited cells with strong nuclear YAP (Fig. 6c).

We also found an opposing correlation between high YAP expression and patient survival in luminal (ER+, PR+) breast cancer and basal (triple-negative) (HER2-, ER-, PR-) breast cancer, as shown by Kaplan Meier analyses (Fig. 6d, e)^69^. The high expression of YAP in luminal breast cancer correlates with an increase in patient survival, while it correlated with a decrease in survival in triple negative breast cancer patients. The high expression of YAP target genes such as *CTGF, CYR61 and IGFBP3* correlated with reduced survival of patients with triple-negative breast cancer (Fig. 6f). Overall, these data show that YAP is highly expressed in basal (triple-negative) breast cancers, and its quantification can be used to predict patient survival in a subtype-dependent manner.

## Discussion

Here we examined the essential role of the transcriptional co-activator YAP in basal breast cancer. We identified a key mechanism demonstrating the oncogenic co-operation of Met, YAP and β-catenin in basal breast cancer (see schemes in Fig. 7). We show that Met signalling controls the nuclear translocation and activation of YAP and β-catenin. Upon translocation, YAP and β-catenin interact and bind to enhancer regions of Wnt target genes leading to gene transcription. In the absence of YAP, Met signalling can no longer induce the nuclear translocation of β-catenin, sequestering it in the cytoplasm and preventing the transcription of Wnt target genes. Thus, YAP is a key bottleneck required for the oncogenic activity of Met and β-catenin in basal breast cancer.

**Figure 7.**
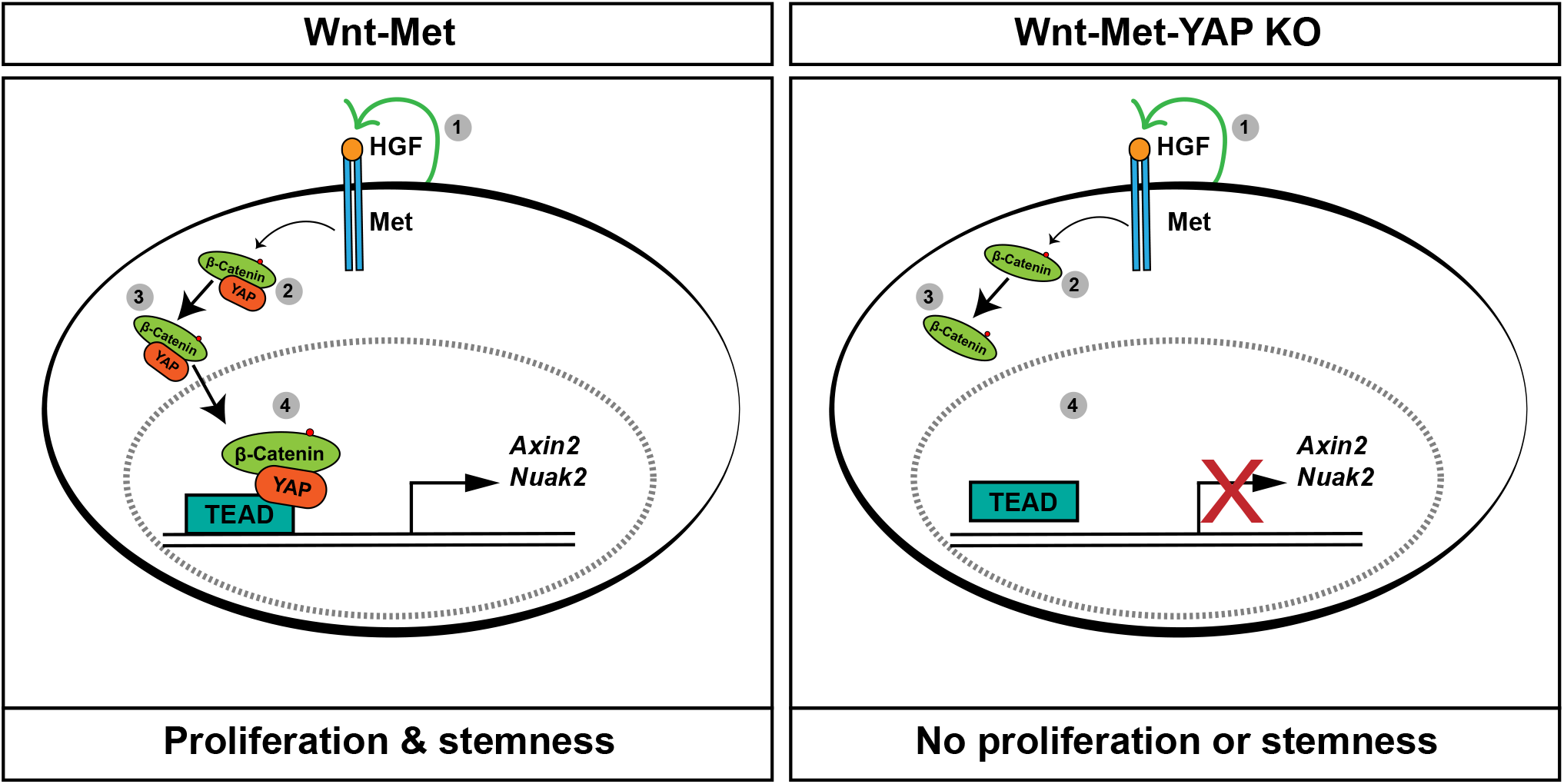
Schematic model for HGF/Met regulation of β-catenin and YAP activity. **a.** (1) Extracellular HGF binds the Met receptor leading to hetero-dimerization and intracellular tyrosine phosphorylation. (2) β-catenin and YAP are activated and translocated to the nucleus, (3) where they interact and bind the transcriptional co-activator, TEAD leading to the transcription of target genes. **b.** (1) HGF binds the Met receptor leading to hetero-dimerization and tyrosine phosphorylation. (2) In the absence of YAP, β-catenin is sequestered in the cytoplasm, (3) and can no longer induce the transcription of genes such as *Axin2*.

YAP activity has been shown to be regulated by several upstream mechanisms^10,14,71,70^. Utilizing our genetic mouse model, proteomic and phosphor-proteomic approaches, we found that Met signalling promotes YAP activity. We identified two phosphorylation sites that may aid YAP activation. Moreover, our analysis revealed that the regulation of basal characteristics by Met depends on YAP. This suggests that YAP might also play a major role in the de-differentiation process of other models of basal breast cancers^72^. Our findings demonstrate that Met signalling confers its oncogenic and metastatic abilities through activating YAP. It would be interesting to investigate if bypassing the Met receptor by overexpression of only YAP and β-catenin lead to tumours that are identical to that of Wnt-Met.

Wnt-Met activation generates cancer-propagating cells^11^. Wnt/β-catenin activity is known to be enhanced in CSCs and confers breast cancer re-occurrence and progression^73,74^. Several studies have suggested that Wnt/β-catenin is linked to Hippo/YAP activity^30,63,75^. In the present study, we found both YAP and β-catenin target genes were increased in Wnt-Met tumours. Moreover, we identified active YAP in the CSCs of these tumours. Functionally, we show that YAP controls CSC properties: through pharmacological and genetic knockouts of YAP, we demonstrated that the absence of YAP strongly delayed Wnt-Met-driven tumorigenesis and decreased the expansion of CD24^hi^CD49f^hi^ CSCs. Furthermore, we established that Wnt/β-catenin target gene expression was decreased upon YAP inhibition, demonstrating that YAP is required for Wnt/β-catenin-dependent CSC expansion in Wnt-Met tumours.

Wnt and Hippo signalling have been described as tightly intertwined. For example, in the mouse liver, the co-activation of β-catenin and YAP have been shown to lead to the rapid generation of hepatocellular carcinoma^76^. It has been shown that the absence of Wnt signals results in YAP/TAZ sequestration in the cytoplasm by the β-catenin destruction complex, comprised of APC, Axin and GSK3^30^. In this context, the knockout of YAP/TAZ in embryonic stem cells leads to the activation of β-catenin and compensates for loss of Wnt signalling. However, we found that genetic ablation of YAP controls β-catenin activity at two levels: i) in the nucleus through co-binding the regulatory regions of β-catenin target genes such as *Axin2*, and ii) by regulating β-catenin nuclear translocation, showing that basal breast cancers require YAP for nuclear β-catenin activity. It is likely that the master regulatory control of Wnt target gene expression by YAP is both context- and tissue-dependent.

YAP’s relevance in breast cancer is currently under dispute. YAP was recently shown to be inactive and even downregulated in ductal carcinoma *in situ* (DCIS)^38^. In addition, breast cancer patients displaying high YAP activity appeared to have a higher overall positive prognosis, leading to the belief that YAP might act as a tumour suppressor in breast cancer^37,77^. In this scenario, keeping YAP activity under tight control is thought to help tumour cells escape immunosurveillance^78^. However, other oncogenic transcriptional systems are equally likely to stimulate a strong immune response in the host^79,80^. These studies did not investigate the functional role of YAP in a context-dependent manner such as breast cancer or tumour cell subtypes. Our study addresses these issues by using genetic, siRNA and pharmacological interference with YAP in basal breast cancer to demonstrate its function specifically in CSCs. In basal breast cancer, we show that YAP’s genetic ablation strongly delayed Wnt-Met-driven tumorigenesis. Segregation of luminal (ER+, PR+) breast cancer patients and basal, triple-negative patients revealed that YAP expression correlated with patient survival in opposing ways. High YAP activity was associated with prolonged survival in luminal breast cancer patients, but decreased survival in basal/triple-negative patients. Thus, YAP appears to function as an oncogene in some cancer subtypes and a tumour suppressor gene in others. Further studies will be required to understand what mechanisms are responsible for the context-dependent function of YAP and whether its inhibition in breast cancer has converse therapeutic benefits in basal and luminal breast cancers.

In conclusion, our data demonstrate that YAP is essential for the co-operation of Wnt and RTK (Met) signalling in initiating and maintaining basal mammary gland tumours. We demonstrate that Met signalling controls the de-differentiation of luminal to basal mammary gland cells through the downstream activation of YAP. Furthermore, we show that YAP is highly expressed in the CSCs of Wnt-Met mice and promotes their expansion, partly by regulating the activity of β-catenin. We confirmed elevated YAP activity in mammospheres of human basal breast cancer cell lines, in tumours from human patients with basal breast cancer, and in PDX models of basal breast cancer and contrasted these findings to the luminal stages. Our data highlight the importance of YAP as a crucial tumour initiator and CSC regulator in basal breast cancer and demonstrate that its activity has prognostic value when comparing basal with luminal breast cancers. This suggests that targeting YAP through specific new drugs is a potential therapeutic avenue for treating basal breast cancers in the future.

## Acknowledgements

We thank Russ Hodge of the MDC for improving our manuscript style, Dr. Hans-Peter Rahn and the FACS Core Facility of the MDC for their expertise, Dr. Anca Margineanu and the Confocal Microscopy Core Facility of the MDC for the support with microscopy, and Dr. Diana Behrens at EPO-Buch for supervising the *in vivo* drug treatments.

## Author Contributions

H.Q and W.B conceived the project. H.Q designed, carried out and interpreted experiments. R.V provided technical assistance, acquired data and managed animal experiments. O.P and P.M carried out and analysed proteomic and phospho-proteomic experiments and data. E. M provided PDX material. C.M analysed chip-seq data. H.Q and W.B wrote the paper with contributions from E.K and Y.F. W.B and Y.F supervised the project.

## Methods

### Mice

All animal experiments were conducted in accordance with European, National and MDC regulations. Wnt-Met mutant mice were previously described^18,24^. Yap1tm1.1Dupa/J flox mice were purchased from Jackson Laboratories (Stock No: 027929). Animal experiments were approved by the Ethical Board of the ‘Landesamt für Gesundheit und Soziales (LaGeSo)’, Berlin. Mice were induced via pregnancy between 8 - 12 weeks old. Tumours were harvested between 1 and 2 weeks post-partum and when they reached a maximum size of 1cm^3^.

### Isolation of Mammary Gland Cells

Tumours were minced and digested in DMEM/F12 HAM (Invitrogen) supplemented with 5% FBS (Invitrogen), 5μg/ml insulin (Sigma-Aldrich), 0.5μg/ml hydrocortisone (Sigma-Aldrich), 10ng/ml EGF (Sigma-Aldrich) (Digestion medium) containing 300 U/ml Collagenase type III (Worthington), 100U/ml Hyaluronidase (Worthington) and 20μg/ml Liberase TM (Sigma) at 37°C for 1.5 hours shaking. Resulting organoids were re-suspended in 0.25% trypsin-EDTA (Invitrogen) at 37°C for 1 minute and further dissociated in digestion medium containing 2 mg/ml Dispase (Invitrogen) and 0.1 mg/ml DNase I (Worthington) at 37°C for 45mins while shaking. Samples were filtered with 40 μm cell strainers (BD Biosciences) and incubated with 0.8% NH4Cl solution on ice for 3 minutes (RBC lysis). Lysis was stopped by washing in 30ml DPBS w/o Ca^2+,^ Mg^2+^ (Gibco, Cat. #14190169). Resulting pellets were used for downstream applications outlined below.

### Organotypic Stem Cell-enriched 3D Cultures

Organotypic stem cell-enriched cultures were generated using previously published protocols^57,58^ with minor adjustments. Single cells from digested mammary glands were resuspended and plated on Collagen I-coated plates (50μg/ml) in stem cell-enriching medium MEBM (Lonza Cat. #CC-3151), supplemented with 2% B27 (Invitrogen, Cat. # 17504044), 20 ng/ml bFGF (Invitrogen, Cat. # 13256029), 20 ng/ml EGF (Sigma, Cat. #SRP3196-500μg), 4 μg/ml heparin (Sigma, Cat. # H3149), 5 μg/ml insulin (Sigma, Cat. #I0516-5ml), 0.5μg/ml hydrocortisone (Sigma, Cat. #H0888-1G) and 1X Gentamicin (Sigma, Cat. # G1397-100ML) for 16-18 hours. Cells were washed twice with DPBS, washed with 0.25% trypsin-EDTA for 30 seconds and then incubated with 0.25% trypsin-EDTA at 37°C for 5 - 7 minutes until cells detached. Trypsin was inactivated with DMEM/F12 supplemented with 10% FBS, 1% HEPES and 1% Pen/Strep. Cells were re-suspended in stem-enriching medium and counted. Cells were then seeded in 25μl droplets containing 50% reduced growth factor Matrigel at a density of 100cells/μl. Plates were carefully flipped and Matrigel was let solidify at 37°C for 45mins - 1hour. 0.5ml of stem medium was added per well of 24-well plates containing a 25μl droplet. Medium was changed every second day. For secondary sphere formation, spheres were dissociated for 10mins in 0.25% trypsin-EDTA, and cells were filtered using 0.45μm filters. Single cell suspensions were then re-seeded as described for primary cells.

### Fluorescence-Activated Cell Sorting (FACS)

Single cell suspensions from dissociated tumours were re-suspended at 10,000 cells/μl and incubated with conjugated primary antibodies at 4°C for 15mins. Cells were then washed x3 in DPBS and incubated with 7AAD (5μl/10^6^ cells) at room temperature for 5 minutes to stain dead/dying cells. Cells were sorted using the FACSAria II or III (BD Biosciences) or analysed using LSRFortessa (BD Biosciences). Compensation and unstained controls were carried out for every FACS experiment as required. Data were analysed using FLowJo Analysis Software.

### Histology and Immunostaining

Mammary glands were fixed overnight at 4°C in 4% formaldehyde (Roth). Glands were then dehydrated and embedded in paraffin. For staining, tissue sections were de-paraffinised, rehydrated and stained with Hematoxylin and Eosin (H&E), dehydrated and mounted. Images were acquired with a bright-field Zeiss microscope. For immune-staining, antigen-retrieval was conducted on paraffin-embedded sections by boiling in Tris-EDTA buffer (10mmol/L Tris, 1mmol/L EDTA, pH 9.0) for 20 minutes. Cryo-sections were incubated for 5 minutes at room temperature and then fixed with 4% methanol-free paraformaldehyde for 15 minutes. Adherent cells were fixed with 4% methanol-free paraformaldehyde for 15mins. Stem cell-enriched spheres were collected, centrifuged and then fixed in 4% formaldehyde before being dehydrated and paraffin-embedded. All samples were blocked with 10% horse serum for 1 hour at room temperature prior to being incubated overnight with primary antibodies at 4°C (See Supplementary Table for antibody list). For fluorescent staining, samples were incubated with secondary antibodies and DAPI for 1 hour at room temperature before being mounted with Immu-mount (Thermo Scientific).

For immunohistochemistry, after antigen-retrieval, slides were washed and incubated with 3% H_2_O_2_ for 15 minutes at room temperature, washed 3 times for 5 minutes in PBS and incubated with primary antibodies overnight at 4°C. Horseradish Peroxidase-labelled secondary antibodies were diluted and incubated on slides for 1 hour at room temperature. Slides were then washed and developed using DAB chromogenic substrate (DAKO), dehydrated and mounted.

### Mammary Gland Whole-mount Staining

Mammary glands were dissected, fixed in Carnoy’s fixative for 2 hours, stained in carmine alum solution overnight, de-stained in 70% ethanol, washed in 100% ethanol, defatted in xylene overnight, and mounted using Permount (Thermo Scientific).

### RNA Isolation, cDNA Generation and qPCR

RNA was extracted using TRIzol (Invitrogen). For 3D cultures, TRIzol was added directly to Matrigel. For tissue, snap frozen glands were pulverised on dry ice, and TRIzol was added to 30mg tissue powder. For cell lines, TRIzol was added directly to plate. For mammospheres, spheres were collected, pelleted and TRIzol was added. After 5 minutes at room temperature, 200μl of chloroform was added per 1ml of TRIzol and shook vigorously by hand for 15 seconds. Samples were then centrifuged at 12,000G for 15 minutes. The upper phase was transferred to a fresh Eppendorf tube, and 500μl of isopropanol was added, mixed and incubated at room temperature for 10 minutes. Samples were then centrifuged at 12,000G for 10 minutes. The RNA pellet was washed twice with 1ml of 70% EtOH. All steps were carried out at 4°C unless otherwise stated.

cDNA was generated using MMLV reverse transcriptase (Promega). Real time PCR was performed using Power Up SybR Green master mix (Thermo Scientific #A25741), see Supplementary Table for primer list. GAPDH or RPLO were used as housekeeping genes, to which values were normalised to. Bio-Rad CX manager was used to calculate Cq and analysed data.

### Proteomics and Phospho-proteomics

For proteomics and phospho-proteomics analysis, sample preparation was carried out essentially according to Mertins and collegues, 2018^81^. Briefly, samples were lysed in urea lysis buffer containing protease and phosphatase inhibitors. From each sample, 500 μg was subjected to reduction with DTT and alkylation with iodoacetamide before performing digestion with LysC and trypsin. After desalting, peptides were labelled with TMT6 reagents (Thermo) and fractionated by basic reversed-phase separation into 24 fractions. 1.5 μg per fraction was injected into LC-MS for global proteome analysis, while the remaining majority of the samples were pooled into 12 fractions and enriched for phosphopeptides using robot-assisted iron-based IMAC on an AssayMap Bravo system (Agilent).

Proteomics samples were measured on an Exploris 480 orbitrap mass spectrometer (Thermo) connected to an EASY-nLC system (Thermo). HPLC-separation occurred on an in-house prepared nano-LC column (0.074 mm × 250 mm, 1.9 μm Reprosil C18, Dr Maisch GmbH) using a flow rate of 250 nl/min on a 110 min gradient with an acetonitrile concentration ramp from 4.7% to 55.2% (v/v) in 0.1% (v/v) formic acid. MS acquisition was performed at a resolution of 60,000 in the scan range from 375 to 1,500 m/z. Data-dependent MS2 scans were carried out at a resolution of 45,000 or 30,000 with an isolation window of 0.4 m/z and a maximum injection time of 86 ms (120 ms for phosphoproteomics) using the TopSpeed setting with a 1 sec cycle time. Dynamic exclusion was set to 20 s and the normalised collision energy was specified to 35.

For analysis, the MaxQuant software package version 1.6.10.43^82,83^ was used. TMT6 reporter ion quantitation was turned on using a PIF setting of 0.5. Variable modifications included Met-oxidation, acetylated N-termini and deamidation of Asn and Gln for proteomics analysis. For phosphoproteomics analysis, deamidation was replaced by phosphorylation on Ser, Thr and Tyr. An FDR of 0.01 was applied for peptides, sites and proteins, and the Andromeda search was performed using a mouse Uniprot database (July 2018, including isoforms). Further analysis was done using R and the Protigy package (https://github.com/broadinstitute/protigy). Protein groups were filtered for proteins that have been identified by at least one unique peptide and that had valid TMT reporter ion intensities across all samples. The phosphoproteome was also filtered for sites containing valid values for all samples. For both analyses, two-sample moderated t-tests (limma package^84^) comparing the two groups were calculated using the log2-transformed and median-MAD normalised TMT6 corrected reporter ion intensities, followed by a Benjamini-Hochberg (BH) p-value correction. Heatmaps of selected significant proteins or phosphosites according to BH-adjusted p-value were generated using the pheatmap package^85^ employing hierarchical clustering based on Euclidean distance. For annotation of gene lists derived from a human proteome, i.e. Zanconato and collegues (2016), genes were converted into mouse orthologs using biomaRt^86^ and manual curation.

### Nanostring

PanCancer Pathways panel was purchased from Nanostring including additional custom gene set for selected YAP and β-catenin target genes. 70ng of RNA was hybridised for 16hours at 65°C. Hybridised RNA was loaded onto and analysed using nCounter SPRINT Profiler. Data were analysed using nSolver software. Heatmaps were generated using nSolver software from normalised data using Pearson Correlation. Genes included were manually filtered using available gene lists^40,45^.

### Protein extraction, Western blot and co-IP

For Western blotting, protein was extracted using RIPA buffer with added cOMPLETE™ Mini protease inhibitor cocktail (#11836153001) and phosphatase inhibitor cocktail 2 and 3 (Sigma). For 2D cell culture, cells were scraped into RIPA buffer (150mM NaCl, 1% Triton-X-100, 0.5% sodium deoxycholate, 0.1% SDS, 50mM Tris pH 8.0 and cOMPLETE™ Mini protease inhibitor cocktail (#11836153001) and phosphatase inhibitor cocktail 2 and 3 (Sigma) for lysis. Lysis was carried out on ice for 20mins, samples were then centrifuged at full speed and supernatant was transferred to fresh tubes, and protein concentration was measured using BioRad Bradford reagent.

For co-IP, cells were lysed as above in co-IP buffer (150mM NaCl, 50mM Tris pH 7.5, 1% IGPAL-CA-630 (Sigma #I8896), 5% glycerol, 0.5% deoxycholate, 0.1% SDS with cOMPLETE™ Mini protease inhibitor cocktail (#11836153001) and phosphatase inhibitor cocktail 2 and 3 (Sigma)). Equal amounts of protein were incubated with primary antibodies overnight at 4°C rotating. Immuno-complexes were then captured on Dynabeads Protein G Invitrogen Cat#10003D for 1 hour at 4°C rotating. Samples were washed and boiled in sample buffer and resolved on 10% SDS-PAGE or gradient gels (4 – 20%, Bio-Rad) and electro-transferred to PDVF membrane. Membranes were blocked with 5% BSA for 1 hour and incubated with primary antibodies (see antibody list).

### siRNA

siRNAs were purchased from Thermo Scientific (see Supplementary Table for list of specific siRNAs). 75pmol/well (6-well) of siRNA was transfected using Lipofectamine 2000 following manufacturer’s instructions. Cells were harvested 48 hours post transfection.

### shRNA

PLKO.1 shRNA plasmids were a gift from Yaron Fuchs^32^. Lentiviral packaging plasmids, pMD2.G and psPAX2 were co-transfected along with shscramble and shYAP1 into HEK293 cells. Viruses were collected and concentrated 24 - 48 hours post-transfection. Concentrated viruses were used to transduce Wnt-Met primary mammary cells before being seeded as stem cell-enriched spheres.

### Cell Culture

Breast cancer cell lines were purchased from ATCC (MCF10A, MCF7, BT474, T47D, MDA-MB-231 and BT-549) and Asterand Bioscience (SUM1315 and SUM149). MCF10A cells were maintained in DMEM/F12 Glutamax, 5% horse serum, 20ng/ml EGF, 0.5mg/ml hydrocortisone, 100ng/ml cholera toxin, 10μg/ml insulin and 1% pen/strep. MCF7, BT474 and T47D were maintained in DMEM, 10% FBS, 5μg/ml insulin and 1% pen/strep. MDA-MB-231 and BT-549 were maintained in DMEM, 10% FBS, 1% NEAAs and 1% pen/strep. SUM1315 were maintained in DMEM/F12 HAM-Glutamax, 1% Hepes, 5% FBS, 10ng/ml EGF, 5μg/ml insulin and 1% pen/strep. SUM149 were grown in DMEM/F12 HAM-Glutamax, 1% Hepes, 5% FBS, 1μg/ml hydrocortisone, 5μg/ml insulin and 1% pen/strep. For mammosphere assays, cells were seeded at 5,000cells/ml on poly-HEMA-coated nonadherent 10cm plates in DMEMF12, FGF 10ng/ml, EGF 20ng/ml, ITS 1x and B27 1x. Spheres were grown for up to 10 days with medium supplemented every second day.

### CellTitre-Glo assay

3D cultures were seeded into 96-well opaque-walled plates. Control and treated spheres with 100μl of medium per well were incubated for 30mins at room temperature. 100μl of CellTiter-Glo reagent was added to each well and incubated for 2 minutes shaking to induce cell lysis. Plates were then rested for 10 minutes at room temperature to stabilise the luminescent signal. Luminescence was recorded on a luminometer.

### *In vivo* Inhibitor Studies

For simvastatin treatments, Wnt-Met mice received 100mg/kg of simvastatin or vehicle control via oral gavage daily. Simvastatin was diluted in 2% DMSO, 30% PEG 300, 5% Tween80 and ddH2O. For verteporfin treatments, mice were treated with 2.5mg/kg daily via subcutaneous injection. Verteporfin was diluted in DMSO and then brought to 2.5% DMSO in PBS. Tumour volumes and body weight were determined several times per week.

### PDX models

PDX models were established from triple-negative breast cancer patients with their informed consent as previously described^68^. The experimental protocol and animal housing were in accordance with institutional guidelines as proposed by the French Ethics Committee (Agreement N° B75-05-18).

### ChIP-Atlas data analysis

ChIP-seq tracks for CTNNB1 (SRX833403), TEAD4 (SRX190301) and YAP1 (SRX2844314) in H1-hESC were downloaded from chip-atlas.org^64,87^. Functional annotation for genome regions was taken from ChromHMM^88^. Data were loaded into UCSC genome browser^89^ to make the plots.

### Kaplan-Meier and GOBO analysis

Relapse-free survival of breast cancer patients was analysed using Kaplan-Meier Plotter online software (http://kmplot.com). Trichotomization was set to Q1 vs Q4. Expression analysis of YAP in breast cancer cell lines and human datasets were generated using GOBO online software (http://co.bmc.lu.se/gobo).

## Supplementary Figure Legends

**Supplementary Figure 1.**
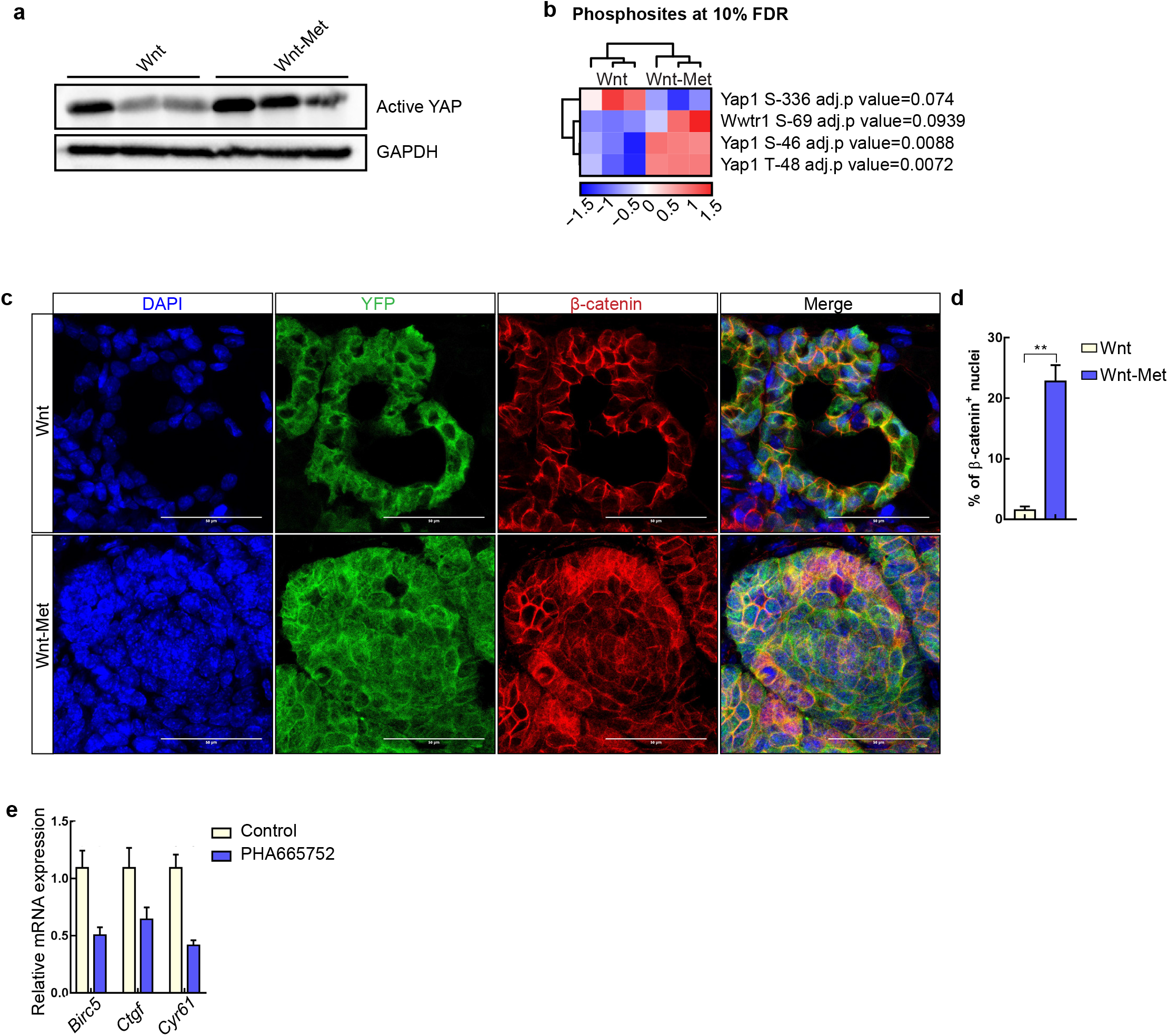
Met signalling regulates YAP and β-catenin. **a.** Western blot for active YAP in triplicates of Wnt and Wnt-Met mammary glands at 1 week PP. **b.** Heatmap of proteomics analysis showing YAP and TAZ (Wwtr1) phosphorylation sites that were significant in a two-sample moderated t-test (adj. p-value ≤ 0.1) revealing significant increase in YAP phosphorylation at S46 and T48 (row-scaling was applied). The heatmap shows median-MAD-normalised input data across all proteins and row-scaling across all samples as was clustered based on Euclidian distance. **c.** Immunofluorescence of YFP (green) and β-catenin (red) in WAPicre; β-cat^GOF^; ROSA26^EYFP^ (Wnt) and WAPicre; Wap-HGF; β-cat^GOF^; ROSA26^EYFP^ (Wnt-Met) at 1-week PP, scale bar, 20μm. **d.** Quantification of β-catenin-positive nuclei (β-catenin^+^ nuclei/number of nuclei per field x 100) in Wnt vs. Wnt-Met tissues. Data shown are mean ±SEM, n=3 biological replicates, **p<0.01, by Student’s t test. **e.** RT-qPCR of YAP target genes in PHA665752-treated mammospheres. Data shown are mean ±SEM, n=2 biological replicates.

**Supplementary Figure 2.**
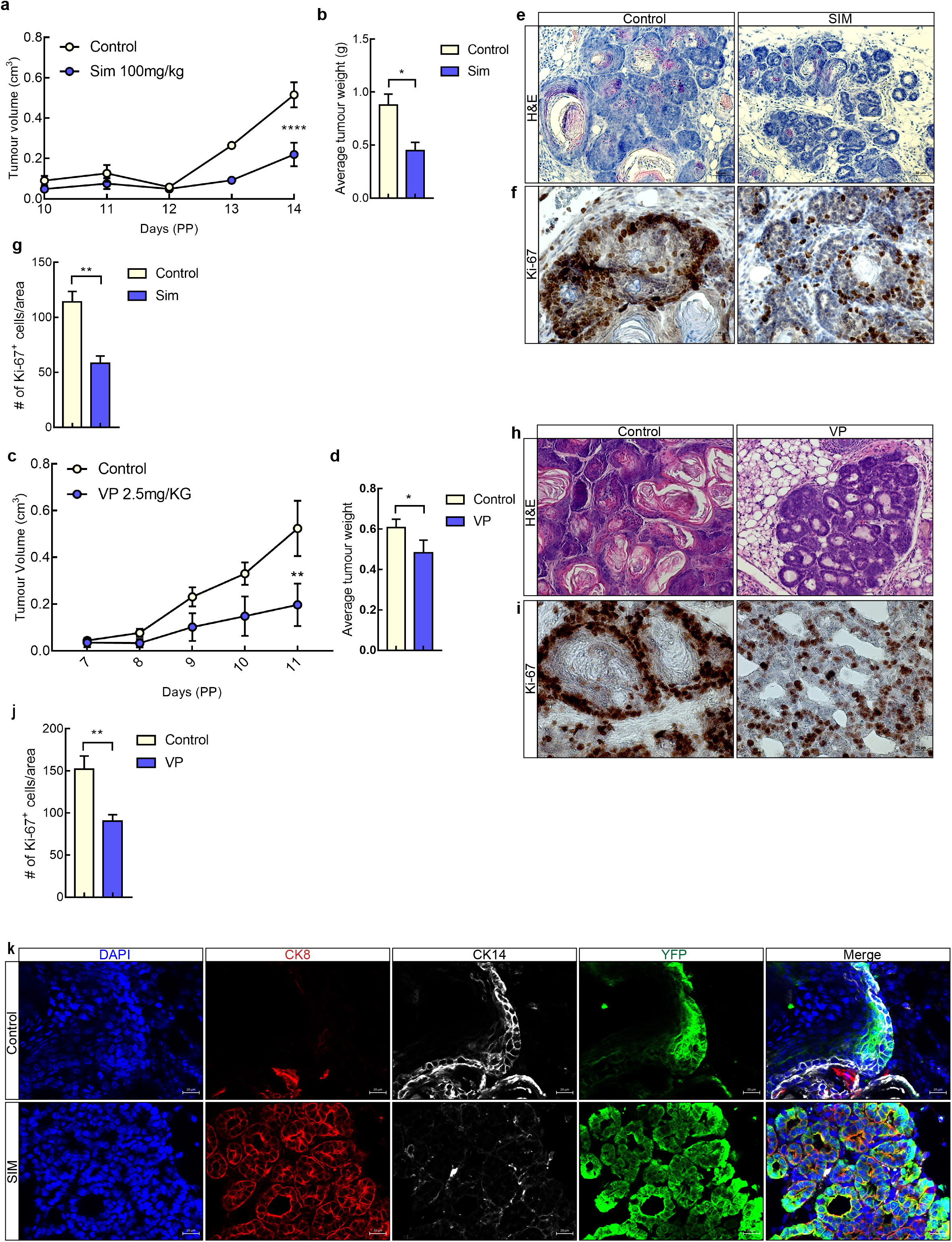
Pharmacological evidence of YAP dependency in Wnt-Met tumours. **a.** Tumour volume curve of Wnt-Met tumours treated with vehicle control or 100mg/kg simvastatin (SIM). Data are mean ±SEM, n=3 biological replicates, ****p<0.0001, Two-way ANOVA, Sidak’s multiple comparisons test. **b.** Average tumour weight of control and SIM-treated mice. Data shown are mean ±SEM, n=3 biological replicates, *p<0.05, by Student’s t test. **c.** Tumour volume curve of Wnt-Met tumours treated with vehicle control or 2.5mg/kg verteporfin (VP). Data are mean ±SEM, n=3 biological replicates, **p<0.01, Twoway ANOVA, Sidak’s multiple comparisons test **d.** Average tumour weight of control and VP treated mice. Data shown are mean ±SEM, n=3 biological replicates, *p<0.05, by Student’s t test. **e.** H&E staining of control and SIM treated Wnt-Met tumours, scale bar, 50μm. **f.** Immunohistochemistry of Ki-67 in control and SIM-treated tumours, scale bar, 20μm. **g.** Quantification of Ki-67-positive cells/area in control and SIM-treated tumours. Data shown are mean ±SEM, n=3 biological replicates, **p<0.01, by Student’s t test. **h.** H&E staining of control and VP-treated Wnt-Met tumours, scale bar, 50μm. **i.** Immunohistochemistry of Ki-67 in control and SIM-treated tumours, scale bar, 20μm. **j.** Quantification of Ki-67-positive cells/area in control and SIM-treated tumours. Data shown are mean ±SEM, n=3 biological replicates, **p<0.01, by Student’s t test. **k.** Immunofluorescent images of control and SIM-treated tumours for CK8 (red), CK14 (white) and YFP (green), scale bar, 20μm.

**Supplementary Figure 3.**
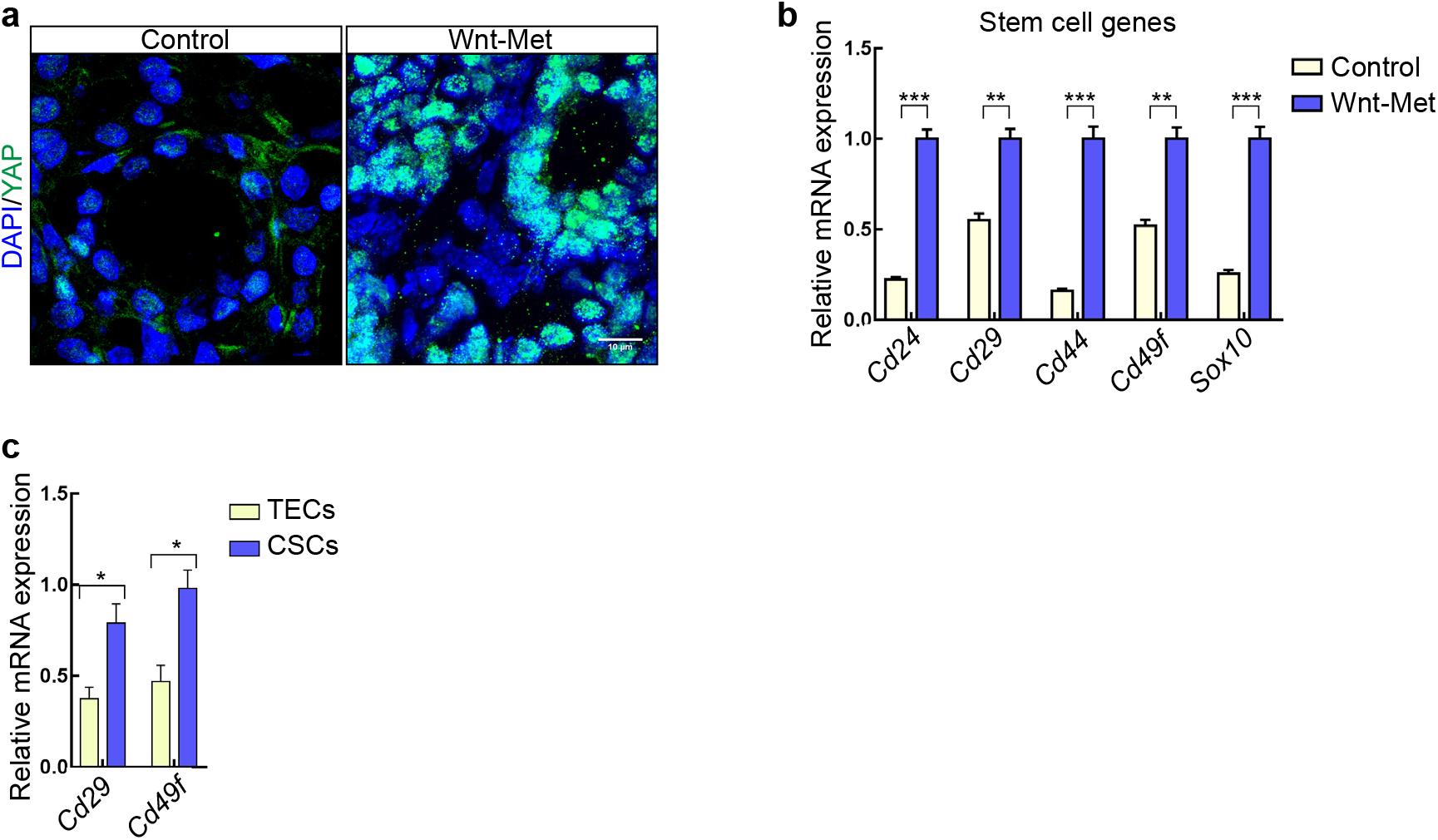
YAP is activated in CSCs of Wnt-Met-driven mammary gland tumours. **a.** Confocal analysis of YAP in control and Wnt-Met tumours, scale bar, 10μm. **b.** qRT-PCR of stem cell genes in Control and Wnt-Met mammary glands. Data shown are mean ±SEM, n=3 biological replicates, **p<0.01, ***p<0.001, by Student’s t-test. **c.** RT-qPCR of *Cd29* and *Cd49f* in TECs and CSCs. Data shown are mean ±SEM, n=3 biological replicates, *p<0.05, by Student’s t-test.

**Supplementary Figure 4.**
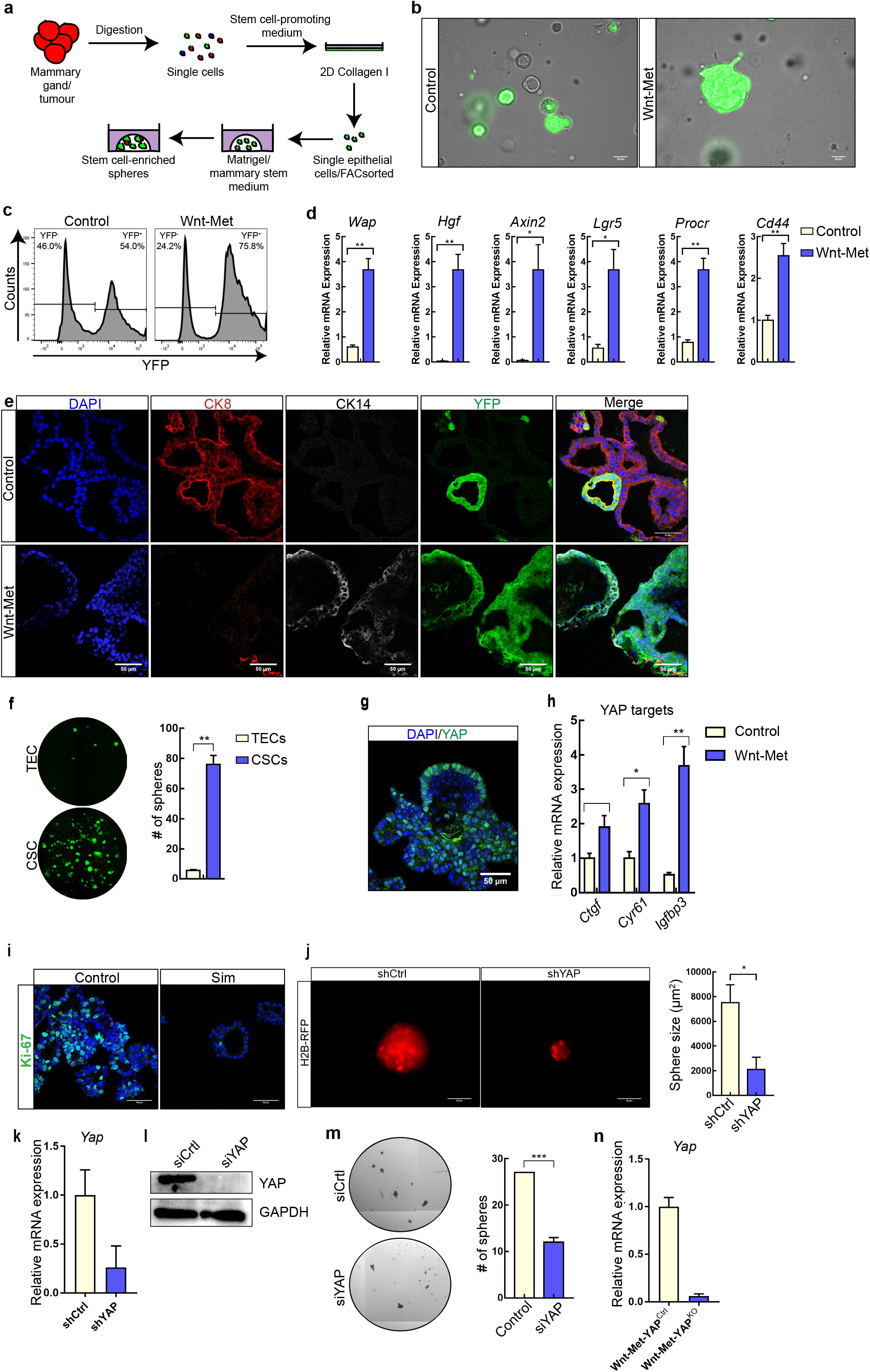
Stem cell-enriched spheres recapitulate in vivo cell phenotypes. **a.** Scheme showing the method used to isolate and generate stem cell-enriched spheres. **b.** Bright-field and fluorescent images of stem cell-enriched spheres generated from control or Wnt-Met mammary glands at 2 weeks PP, scale bar, 50μm. **c.** FACS histogram showing YFP-positive cells in control or Wnt-Met derived stem-enriched spheres. Fold-change calculated by YFP cells in Wnt-Met/YFP cells in control. **d.** RT-qPCR of transgene-associated genes in control or Wnt-Met stem cell-enriched spheres. Data shown are mean ±SEM, n=3 biological replicates, *p<0.05, **p<0.01, by Student’s t test. **e.** Confocal images of CK8 (red), CK14 (white) and YFP (green) in control and Wnt-Met stem cell-enriched spheres, scale bar, 50μm. **f.** Fluorescent images showing YFP in stem cell-enriched spheres generated from FACS-isolated TECs and CSCs. Quantification of sphere numbers (right). Data shown are mean ±SEM, n=3 replicates, **p<0.01, by Student’s t-test. **g.** Confocal images of YAP (green) in Wnt-Met stem cell-enriched spheres, scale bar, 50μm. **h.** RT-qPCR YAP target genes in control and Wnt-Met stem cell-enriched spheres. Data shown are mean ±SEM, n=3 biological replicates, *p<0.05, **p<0.01, by Student’s t-test. **i.** Confocal images of Ki-67 (green) in control and 2.5μM simvastatin (SIM)-treated stem cell-enriched spheres, scale bar, 50μm. **j.** Fluorescent images of the H2B-RFP reporter in Wnt-Met stem cell-enriched spheres, scale bar, 50μm. Quantification of sphere numbers on the right. Data shown are mean ±SEM, n=3 biological replicates, *p<0.05, by Student’s t test. **k.** RT-qPCR of *Yap* in sh-treated control (Ctrl) and shYAP Wnt-Met spheres. **l.** Western Blot for YAP in sicontrol and siYAP treated MDA-231 cells, GAPDH was used as a loading control. **m.** Bright-field images of spheres generated from sicontrol and siYAP treated MDA-231 cells. Quantification of sphere numbers on the right. Data shown are mean ±SEM, n=3 replicates ***p<0.001 by Student’s t test. **n.** RT-qPCR of *Yap* in Wnt-Met-YAP^Ctrl^ and Wnt-Met-YAP^KO^ spheres.

**Supplementary Figure 5.**
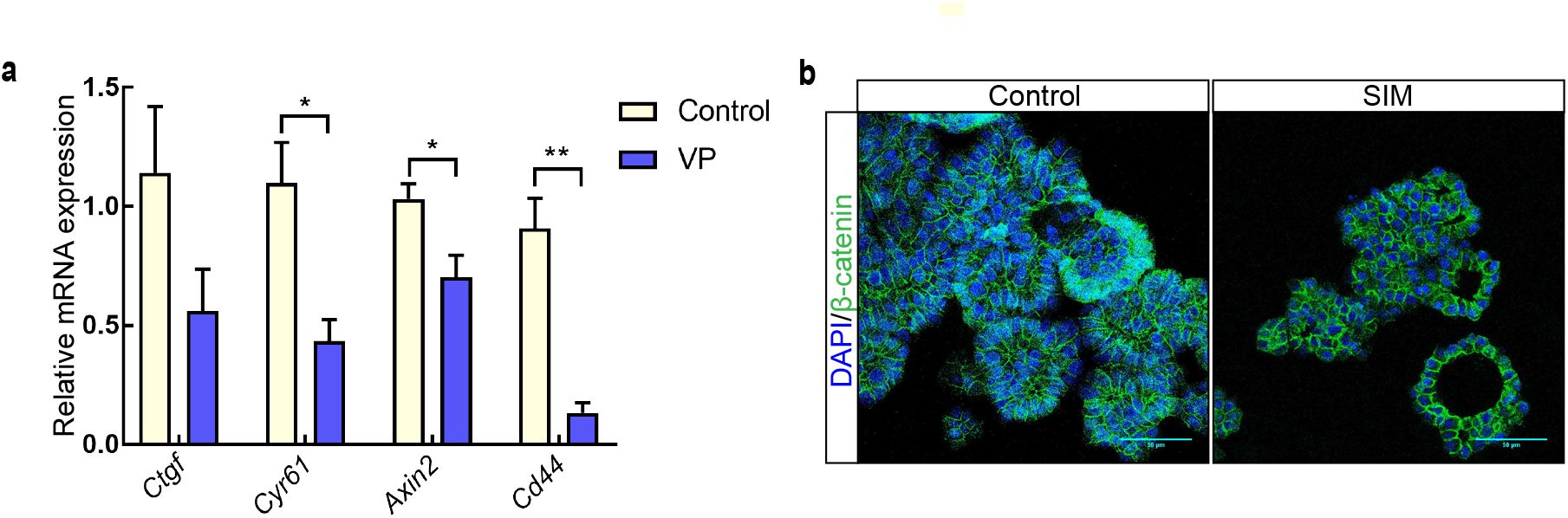
YAP controls β-catenin activity. **a.** RT-qPCR of YAP and β-catenin target genes in Wnt-Met mammary gland spheres treated with 2μM verteporfin (VP) for 48Hrs. Data shown are mean ±SEM, n=3 biological replicates, *p<0.05, **p<0.01, by Student’s t-test. **b.** Confocal images of β-catenin (green) in control and SIM-treated stem cell-enriched spheres, scale bar, 50μm.

**Supplementary Figure 6.**
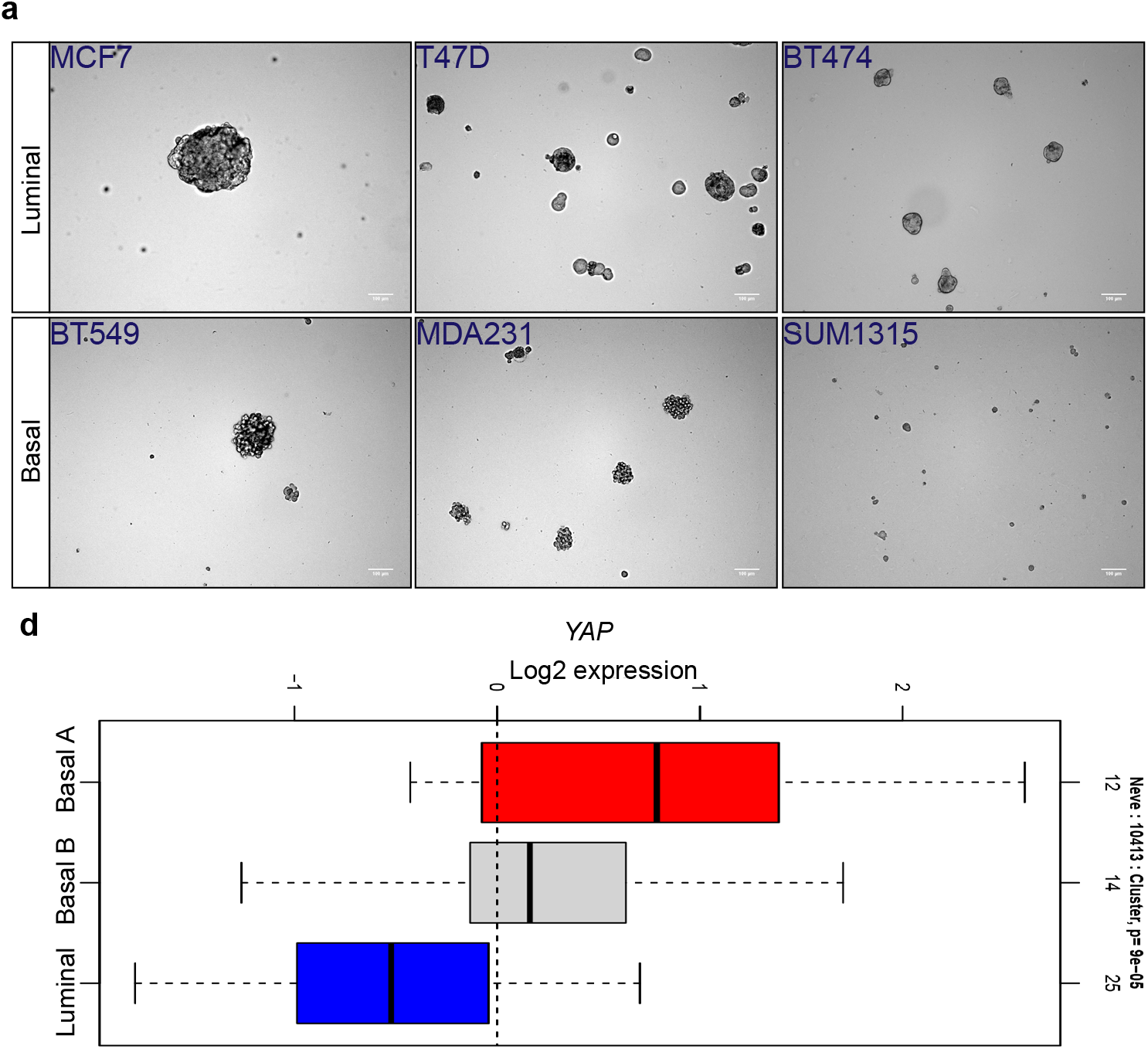
YAP is active in human basal breast tumours and predicts patient outcome in a subtype-dependent manner. **a.** Bright-field images of spheres generated from basal (lower) and luminal (upper) cell lines. Scale bar, 100μm. **b.** Box-plot showing *YAP1* expression in basal and luminal breast tumours.

## References

1. Weigelt, B., Peterse, J. L. & Van’t Veer, L. J. Breast cancer metastasis: Markers and models. Nat. Rev. Cancer 5, 591–602 (2005).

2. Lehmann, B. D. B. et al. Identification of human triple-negative breast cancer subtypes and preclinical models for selection of targeted therapies. J. Clin. Invest. 121, 2750–2767 (2011).

3. Weigelt, B. & Reis-Filho, J. S. Histological and molecular types of breast cancer: is there a unifying taxonomy? Nat. Rev. Clin. Oncol. 6, 718–730 (2009).

4. Carey, L., Winer, E., Viale, G., Cameron, D. & Gianni, L. Triple-negative breast cancer: disease entity or title of convenience? Nat. Rev. Clin. Oncol. 7, 683–692 (2010).

5. Khramtsov, A. I. et al. Wnt / B-Catenin Pathway Activation Is Enriched in Basal-Like Breast Cancers and Predicts Poor Outcome. Am. J. Pathol. 176, 2911–2920 (2010).

6. Koboldt, D. C. et al. Comprehensive molecular portraits of human breast tumours. Nature 490, 61–70 (2012).

7. Gastaldi, S., Comoglio, P. M. & Trusolino, L. The Met oncogene and basal-like breast cancer: another culprit to watch out for? Breast Cancer Res. 12, 208 (2010).

8. Gallego, M. I., Bierie, B. & Hennighausen, L. Targeted expression of HGF / SF in mouse mammary epithelium leads to metastatic adenosquamous carcinomas through the activation of multiple signal transduction pathways. 8498–8508 (2003) doi:10.1038/sj.onc.1207063.

9. Smolen, G. A. et al. Frequent met oncogene amplification in a Brca1/Trp53 mouse model of mammary tumorigenesis. Cancer Res. 66, 3452–3455 (2006).

10. Garcia, S. et al. Poor prognosis in breast carcinomas correlates with increased expression of targetable CD146 and c-Met and with proteomic basal-like phenotype. Hum. Pathol. 38, 830–841 (2007).

11. Holland, J. et al. Combined Wnt/β-Catenin, Met, and CXCL12/CXCR4 Signals Characterize Basal Breast Cancer and Predict Disease Outcome. Cell Rep. 5, 1214–1227 (2013).

12. Khramtsov, A. I. et al. Wnt/beta-catenin pathway activation is enriched in basal-like breast cancers and predicts poor outcome. Am. J. Pathol. 176, 2911–2920 (2010).

13. Lopez-Knowles, E. et al. Cytoplasmic localization of beta-catenin is a marker of poor outcome in breast cancer patients. Cancer Epidemiol. Biomarkers Prev. 19, 301–309 (2010).

14. Gherardi, E., Birchmeier, W., Birchmeier, C. & Woude, G. Vande. Targeting MET in cancer: Rationale and progress. Nat. Rev. Cancer 12, 89–103 (2012).

15. Herschkowitz, J. I. et al. Identification of conserved gene expression features between murine mammary carcinoma models and human breast tumors. Genome Biol. 8, R76 (2007).

16. Shackleton, M. et al. Generation of a functional mammary gland from a single stem cell. Nature 439, 84–88 (2006).

17. Kemper, K., De Goeje, P. L., Peeper, D. S. & Van Amerongen, R. Phenotype switching: Tumor cell plasticity as a resistance mechanism and target for therapy. Cancer Res. 74, 5937–5941 (2014).

18. Hanahan, D. & Weinberg, R. A. Hallmarks of cancer: The next generation. Cell 144, 646–674 (2011).

19. Nassar, D. & Blanpain, C. Cancer Stem Cells: Basic Concepts and Therapeutic Implications. Annu. Rev. Pathol. Mech. Dis. 11, 47–76 (2016).

20. Koren, E. & Fuchs, Y. The bad seed: Cancer stem cells in tumor development and resistance. Drug Resist. Updat. 28, 1–12 (2016).

21. Hallett, R. M., Dvorkin-gheva, A., Bane, A. & Hassell, J. A. CANCER GENOMICS A Gene Signature for Predicting Outcome in Patients with Basal-like Breast Cancer. 17, 1–8 (2012).

22. Mertins, P. et al. Proteogenomics connects somatic mutations to signalling in breast cancer. Nature 534, 55–62 (2016).

23. Barry, E. R. et al. Restriction of intestinal stem cell expansion and the regenerative response by YAP. Nature 493, 106–110 (2013).

24. Qin, H. et al. YAP Induces Human Naive Pluripotency. Cell Rep. 14, 2301–2312 (2016).

25. Rosado-Olivieri, E. A., Anderson, K., Kenty, J. H. & Melton, D. A. YAP inhibition enhances the differentiation of functional stem cell-derived insulin-producing β cells. Nat. Commun. 10, 1–11 (2019).

26. Zhao, B. et al. Inactivation of YAP oncoprotein by the Hippo pathway is involved in cell contact inhibition and tissue growth control. Genes Dev. 21, 2747–2761 (2007).

27. Vassilev, A., Kaneko, K. J., Shu, H., Zhao, Y. & DePamphilis, M. L. TEAD/TEF transcription factors utilize the activation domain of YAP65, a Src/Yes-associated protein localized in the cytoplasm. Genes Dev. 15, 1229–1241 (2001).

28. Zhao, B. et al. TEAD mediates YAP-dependent gene induction and growth control. Genes Dev. 22, 1962–1971 (2008).

29. Xia, H. et al. EGFR-PI3K-PDK1 pathway regulates YAP signaling in hepatocellular carcinoma: The mechanism and its implications in targeted therapy article. Cell Death Dis. 9, (2018).

30. Azzolin, L. et al. YAP / TAZ Incorporation in the b -Catenin Destruction Complex Orchestrates the Wnt Response. Cell 158, 157–170 (2014).

31. Sorrentino, G. et al. Glucocorticoid receptor signalling activates YAP in breast cancer. Nat. Commun. 8, 1–14 (2017).

32. Yosefzon, Y. et al. Caspase-3 Regulates YAP-Dependent Cell Article Caspase-3 Regulates YAP-Dependent Cell Proliferation and Organ Size. Mol. Cell 70, 573–587.e4 (2018).

33. Fu, V., Plouffe, S. W. & Guan, K. L. The Hippo pathway in organ development, homeostasis, and regeneration. Curr. Opin. Cell Biol. 49, 99–107 (2017).

34. Chang, S. S. et al. Aurora A kinase activates YAP signaling in triple-negative breast cancer. Oncogene 36, 1265–1275 (2017).

35. Song, S. et al. Hippo coactivator YAP1 upregulates SOX9 and endows esophageal Cancer cells with stem-like properties. Cancer Res. 74, 4170–4182 (2014).

36. Zanconato, F. et al. Transcriptional addiction in cancer cells is mediated by YAP/TAZ through BRD4. Nat. Med. 24, 1599–1610 (2018).

37. Elster, D. et al. TRPS1 shapes YAP/TEAD-dependent transcription in breast cancer cells. Nat. Commun. 9, (2018).

38. Lee, J. Y. et al. YAP-independent mechanotransduction drives breast cancer progression. Nat. Commun. 1–9 (2019) doi:10.1038/s41467-019-09755-0.

39. Kurppa, K. J. et al. Treatment-Induced Tumor Dormancy through YAP-Mediated Transcriptional Reprogramming of the Apoptotic Pathway. Cancer Cell 37, 104–122.e12 (2020).

40. Zanconato, F. et al. Genome-wide association between YAP/TAZ/TEAD and AP-1 at enhancers drives oncogenic growth. Nat. Cell Biol. 17, 1218–1227 (2015).

41. Barker, N. et al. Identification of stem cells in small intestine and colon by marker gene Lgr5. Nature 449, 1003–1007 (2007).

42. Muramatsu, T. et al. YAP is a candidate oncogene for esophageal squamous cell carcinoma. Carcinogenesis 32, 389–398 (2011).

43. Liao, M. J. et al. Enrichment of a population of mammary gland cells that form mammospheres and have in vivo repopulating activity. Cancer Res. 67, 8131–8138 (2007).

44. Dontu, G. et al. In vitro propagation and transcriptional profiling of human mammary stem/progenitor cells. Genes Dev. 17, 1253–1270 (2003).

45. Herbst, A. et al. Comprehensive analysis of β-catenin target genes in colorectal carcinoma cell lines with deregulated Wnt/β-catenin signaling. BMC Genomics 15, (2014).

46. Zhang, N. et al. Article The Merlin / NF2 Tumor Suppressor Functions through the YAP Oncoprotein to Regulate Tissue Homeostasis in Mammals. Dev. Cell 19, 27–38 (2010).

47. Sorrentino, G. et al. Metabolic control of YAP and TAZ by the mevalonate pathway. Nat. Cell Biol. 16, 357–366 (2014).

48. Liu-chittenden, Y. et al. Genetic and pharmacological disruption of the TEAD–YAP complex suppresses the oncogenic activity of YAP. Genes Dev. 1300–1305 (2012) doi:10.1101/gad.192856.112.

49. Di-Cicco, A. et al. Paracrine met signaling triggers epithelial–mesenchymal transition in mammary luminal progenitors, affecting their fate. Elife 4, 1–25 (2015).

50. Stingl, J. et al. Purification and unique properties of mammary epithelial stem cells. Nature 439, 993–997 (2006).

51. Shipitsin, M. et al. Molecular Definition of Breast Tumor Heterogeneity. Cancer Cell 11, 259–273 (2007).

52. Dravis, C. et al. Sox10 Regulates Stem/Progenitor and Mesenchymal Cell States in Mammary Epithelial Cells. Cell Rep. 12, 2035–2048 (2015).

53. Lustig, B. et al. Negative feedback loop of Wnt signaling through upregulation of conductin/axin2 in colorectal and liver tumors. Langenbeck’s Arch. Surg. 386, 466 (2001).

54. Kim, T. et al. A basal-like breast cancer-specific role for SRF–IL6 in YAP-induced cancer stemness. Nat. Commun. 6, 10186 (2015).

55. Lian, I. et al. The role of YAP transcription coactivator in regulating stem cell selfrenewal and differentiation. Genes Dev. 24, 1106–1118 (2010).

56. Wang, D. et al. Identification of multipotent mammary stem cells by protein C receptor expression. Nature 517, 81–4 (2015).

57. Jechlinger, M., Podsypanina, K. & Varmus, H. Regulation of transgenes in threedimensional cultures of primary mouse mammary cells demonstrates oncogene dependence and identifies cells that survive deinduction. Genes Dev. 23, 1677–1688 (2009).

58. Jechlinger, M. Organotypic culture of untransformed and tumorigenic primary mammary epithelial cells. Cold Spring Harb. Protoc. 2015, 457–461 (2015).

59. Foster, B. M., Zaidi, D., Young, T. R., Mobley, M. E. & Kerr, B. A. CD117/c-kit in cancer stem cell-mediated progression and therapeutic resistance. Biomedicines 6, 1–19 (2018).

60. Oakes, S. R., Gallego-Ortega, D. & Ormandy, C. J. The mammary cellular hierarchy and breast cancer. Cell. Mol. Life Sci. 71, 4301–4324 (2014).

61. Konstantinou, E. K. et al. Verteporfin-induced formation of protein cross-linked oligomers and high molecular weight complexes is mediated by light and leads to cell toxicity. Sci. Rep. 7, 1–11 (2017).

62. Geiss, G. K. et al. Direct multiplexed measurement of gene expression with color-coded probe pairs. Nat. Biotechnol. 26, 317–325 (2008).

63. Rosenbluh, J. et al. b -Catenin-Driven Cancers Require a YAP1 Transcriptional Complex for Survival and Tumorigenesis. Cell 151, 1457–1473 (2012).

64. Oki, S. et al. Ch IP-Atlas: a data-mining suite powered by full integration of public ChIP-seq data. EMBO Rep. 19, 1–10 (2018).

65. Galli, G. G. et al. YAP Drives Growth by Controlling Transcriptional Pause Release from Dynamic Enhancers Short Article YAP Drives Growth by Controlling Transcriptional Pause Release from Dynamic Enhancers. Mol. Cell 60, 328–337 (2015).

66. Yuan, W. C. et al. NUAK2 is a critical YAP target in liver cancer. Nat. Commun. 9, 4834 (2018).

67. Ringnér, M., Fredlund, E., Häkkinen, J., Borg, Å. & Staaf, J. GOBO: Gene expression-based outcome for breast cancer online. PLoS One 6, 1–11 (2011).

68. Coussy, F. et al. A large collection of integrated genomically characterized patient-derived xenografts highlighting the heterogeneity of triple-negative breast cancer. Int. J. Cancer 1912, 1902–1912 (2019).

69. Gyorffy, B. et al. An online survival analysis tool to rapidly assess the effect of 22,277 genes on breast cancer prognosis using microarray data of 1,809 patients. Breast Cancer Res. Treat. 123, 725–731 (2010).

70. Ponzo, M. G. et al. Met induces mammary tumors with diverse histologies and is associated with poor outcome and human basal breast cancer. Proc. Natl. Acad. Sci. U. S. A. 106, 12903–12908 (2009).

71. Azad, T. et al. A gain-of-functional screen identifies the Hippo pathway as a central mediator of receptor tyrosine kinases during tumorigenesis. Oncogene 39, 334–355 (2020).

72. Molyneux, G. et al. Article BRCA1 Basal-like Breast Cancers Originate from Luminal Epithelial Progenitors and Not from Basal Stem Cells. Stem Cell 7, 403–417 (2010).

73. Jang, G. B. et al. Blockade of Wnt/β-catenin signaling suppresses breast cancer metastasis by inhibiting CSC-like phenotype. Sci. Rep. 5, 1–15 (2015).

74. Cleary, A. S., Leonard, T. L., Gestl, S. A. & Gunther, E. J. Tumour cell heterogeneity maintained by cooperating subclones in Wnt-driven mammary cancers. Nature 508, 113–117 (2014).

75. Park, H. W. et al. Alternative Wnt Signaling Activates YAP/TAZ. Cell 162, 780–794 (2015).

76. Tao, J. et al. Activation of β-catenin and Yap1 in human hepatoblastoma and induction of hepatocarcinogenesis in mice. Gastroenterology 147, 690–701 (2014).

77. von Eyss, B. et al. A MYC-Driven Change in Mitochondrial Dynamics Limits YAP/TAZ Function in Mammary Epithelial Cells and Breast Cancer. Cancer Cell 28, 743–757 (2015).

78. Moroishi, T. et al. The Hippo Pathway Kinases LATS1/2 Suppress Cancer Immunity. Cell 167, 1525–1539.e17 (2016).

79. Raulet, D. H. & Guerra, N. Oncogenic stress sensed by the immune system: Role of natural killer cell receptors. Nat. Rev. Immunol. 9, 568–580 (2009).

80. Nepal, R. M. et al. AID and RAG1 do not contribute to lymphomagenesis in Eμ c-myc transgenic mice. Oncogene 27, 4752–4756 (2008).

81. Mertins, P. et al. Reproducible workflow for multiplexed deep-scale proteome and phosphoproteome analysis of tumor tissues by liquid chromatography-mass spectrometry. Nat. Protoc. 13, 1632–1661 (2018).

82. Cox, J. et al. Andromeda: A peptide search engine integrated into the MaxQuant environment. J. Proteome Res. 10, 1794–1805 (2011).

83. Cox, J. & Mann, M. MaxQuant enables high peptide identification rates, individualized p.p.b.-range mass accuracies and proteome-wide protein quantification. Nat. Biotechnol. 26, 1367–1372 (2008).

84. Ritchie, M. E. et al. Limma powers differential expression analyses for RNA-sequencing and microarray studies. Nucleic Acids Res. 43, e47 (2015).

85. Kolde, R. pheatmap: Pretty Heatmaps. R package version 1.0.12. https://CRAN.R-project.org/package=pheatmap (2019).

86. Durinck, S. et al. BioMart and Bioconductor: A powerful link between biological databases and microarray data analysis. Bioinformatics 21, 3439–3440 (2005).

87. Ernst, J. et al. Mapping and analysis of chromatin state dynamics in nine human cell types. Nature 473, 43–49 (2011).

88. Ernst, J. & Kellis, M. ChromHMM: Automating chromatin-state discovery and characterization. Nat. Methods 9, 215–216 (2012).

89. Kent, W. J. et al. The Human Genome Browser at UCSC. Genome Res. 12, 996–1006 (2002).

